# Allosteric mechanism of membrane fusion activation in a herpesvirus

**DOI:** 10.1101/2024.09.20.610514

**Authors:** Gonzalo L. González-Del Pino, Richard M. Walsh, Doina Atanasiu, Tina M. Cairns, Wan Ting Saw, Gary H. Cohen, Ekaterina E. Heldwein

**Affiliations:** Department of Molecular Biology and Microbiology, Tufts University School of Medicine, Boston, MA 02111; Tufts NIH-IRACDA program, Tufts University School of Medicine, Boston, MA 02111; Harvard Medical School Cryoelectron Microscopy Center, Boston, MA 02115; University of Pennsylvania School of Dental Medicine, Philadelphia, PA 19104

**Author notes:** Denotes these authors contributed equally to the work described in this article (Gonzalo L. González-Del Pino, Richard M. Walsh). Corresponding author (EEH).

**Keywords:** herpesvirus, herpes simplex virus, HSV-2, gB, gH/gL, viral entry, membrane fusion, glycoprotein, viral fusogen, cryoEM, antibody, complex, activation, conformation, ensembles, regulation, conformational rearrangement, refolding

## Abstract

*Herpesviridae* infect nearly all humans for life, causing diseases that range from painful to life-threatening^1^. These viruses penetrate cells by employing a complex apparatus composed of separate receptor-binding, signal-transmitting, and membrane-fusing components^2^. But how these components coordinate their functions is unknown. Here, we determined the 4.19-angstrom cryoEM reconstruction of the central signal-transmitting component from herpes simplex virus 2, the gH/gL complex, in its elusive pre-activation state. Analysis of the continuum of conformational ensembles observed in cryoEM data revealed a series of structural rearrangements in gH/gL that allosterically transmit the fusion-triggering signal from the receptor-binding glycoprotein gD to the membrane fusogen gB. Furthermore, we identified a structural “switch” element in gH/gL that refolds and flips 180 degrees during the transition from pre-activation to activated form. Conservation of this “switch” in gH/gL homologs suggests that the proposed fusion triggering mechanism may apply to all *Herpesviridae* and points to a new target for subunit-based vaccines and treatment efforts.

## INTRODUCTION

*Herpesviridae*, commonly known as Herpesviruses, are ancient DNA viruses that infect mammals, birds, and reptiles, and establish lifelong infections in >90% of people worldwide^3^. While often asymptomatic, herpesvirus infections can cause diseases ranging from painful mucocutaneous lesions to loss of vision, encephalitis, cancer, developmental abnormalities, and disseminated disease^3,4^. Current therapeutic and prophylactic options are very limited^5,6^.

Herpesviruses penetrate target host cells by binding an entry receptor on the cell surface and fusing their lipid envelopes with the plasma membrane or membrane of an endocytic vesicle^7^. Unusually, herpesviruses distribute entry functions – host receptor engagement, fusion regulation, and membrane fusion – across several viral glycoproteins^8,9^. Entry of the prototypical herpes simplex viruses 1 and 2 (HSV-1 and HSV-2) requires four viral glycoproteins: gD, gB, gH, and gL^2,9,10^. gD is a receptor-binding protein that engages one of its three cognate receptors: nectin-1^11,12^, a cell adhesion molecule; herpes virus entry mediator (HVEM)^12–14^, an immune modulator; or an O-sulfonated heparan sulfate^10,15,16^. gB is a class III viral fusogen that catalyzes the merger of the viral and host membranes^17,18^. gH and gL form a uniquely folded heterodimeric complex, gH/gL, that somehow “senses” the receptor-binding status of gD and, in response, activates the fusogenic activity of gB^10,19–21^. Indeed, gH/gL can interact with both gD and gB^22^.

Previously, we reported the crystal structure of the N-terminally truncated gH/gL ectodomain from HSV-2, lacking residues 19-48 immediately following the signal sequence (gH_ectoΔ48_/gL)^19^. This structure is thought to represent an activated state of gH/gL because the gH_Δ48_/gL variant supports low-level membrane fusion in the absence of gD, which suggests that the N terminus of gH could be autoinhibitory^23^. Its removal may activate gH/gL by mimicking conformational changes induced by gD/receptor interactions. But how gH/gL transmits the activating signal from gD to gB is unknown.

Here, we report the cryogenic electron microscopy (cryoEM) structure of the HSV-2 gH_ecto_/gL containing intact gH N terminus in its pre-activation state, bound to a fragment of antibody binding (Fab) of a neutralizing antibody CHL27^21^. Comparison with the crystal structure of the gH_ectoΔ48_/gL, representing the activated state, revealed two major structural differences, in addition to changes in domain orientations. First, the N termini of gH and gL, which were unresolved in the activated state, form an X-shaped feature near the gD-binding site that appears to stabilize the pre-activation state. Second, a structural “switch” element near the C terminus of gH refolds and flips towards the gB-binding face of gH/gL. Analysis of the conformational ensembles observed by cryoEM suggests a conformational pathway for refolding of gH/gL from the pre-activation to the activated state, in which interaction of receptor-bound gD (or CHL27 Fab) with the N terminus of gH/gL sets off a series of rearrangements that culminate in the refolding of the C-terminal switch to transmit the activating signal to gB. The proposed allosteric activation mechanism may be conserved across *Herpesviridae*.

## RESULTS

### Structure of the pre-activation state of gH_ecto_/gL

We co-expressed HSV-2 gH_ecto_ with HSV-2 gL in Sf9 cells and purified the heterodimeric complex (**Extended Data Figure 1A and B**). The gH_ecto_/gL retains the gH N terminus, residues 19-48 (immediately following the signal sequence). The N terminus is autoinhibitory^23^. We therefore hypothesized that the gH N terminus may stabilize the pre-activation conformation of gH/gL. The N terminus also contains epitopes of several neutralizing antibodies^21,23,24^. To immobilize this unstructured region, we incubated gH_ecto_/gL with a Fab of CHL27, a mouse neutralizing antibody that binds residues 37-47 and blocks gD binding^21^, and briefly cross-linked the complex with glutaraldehyde. After initial rounds of 3D classification and refinement in cryoSPARC, we obtained an 820,197-particle cryoEM data set. Focused 3D classification and local refinement using a mask focused on the gH/gL portion of the complex yielded a 4.19-Å reconstruction of gH_ecto_/gL from a subset of 223,210 particles that allowed for placement of side chains with reasonable confidence (**Extended Data Figures 2A and B**).

Overall, the structure of the gH_ecto_/gL complex resembles the boot-like structure of the gH_ectoΔ48_/gL (**Figure 1A and Extended Data Figure 3A**). The gH_ecto_ is composed of three domains, H1, H2, and H3 (**Extended Data Figure 3A**). Domain H1 (gH residues 29-332), located in the upper part of the boot, is composed of two subdomains that sandwich the gL core. This N-terminal H1/gL module of gH/gL features a β-sheet, composed of strands from both components, that packs against the helical hairpin of gL (**Extended Data Figure 3B**). The five-β-strand “picket fence” in H1 separates the N-terminal H1/gL module at the top of the boot from the C-terminal module in the lower part of the boot, composed of domains H2 and H3 (**Figure 1A, Extended Data Figure 3A**). The helix-rich domain H2 (gH residues 333-596) in the central part of the boot is composed of a four-helix bundle packed against a six-helix crescent (**Extended Data Figure 3C**). Domain H3, in the toe part of the boot, is a rigid, disc-like β-sandwich surrounded by loops and a helix (**Extended Data Figure 3D**). While the individual domains remain mostly unchanged, the bulk of the heterodimer (H1-gL-H2) is rotated and tilted 8° relative to domain H3 such that the gH_ecto_/gL structure appears to stand more upright (**Figure 1A**).

**Figure 1.**
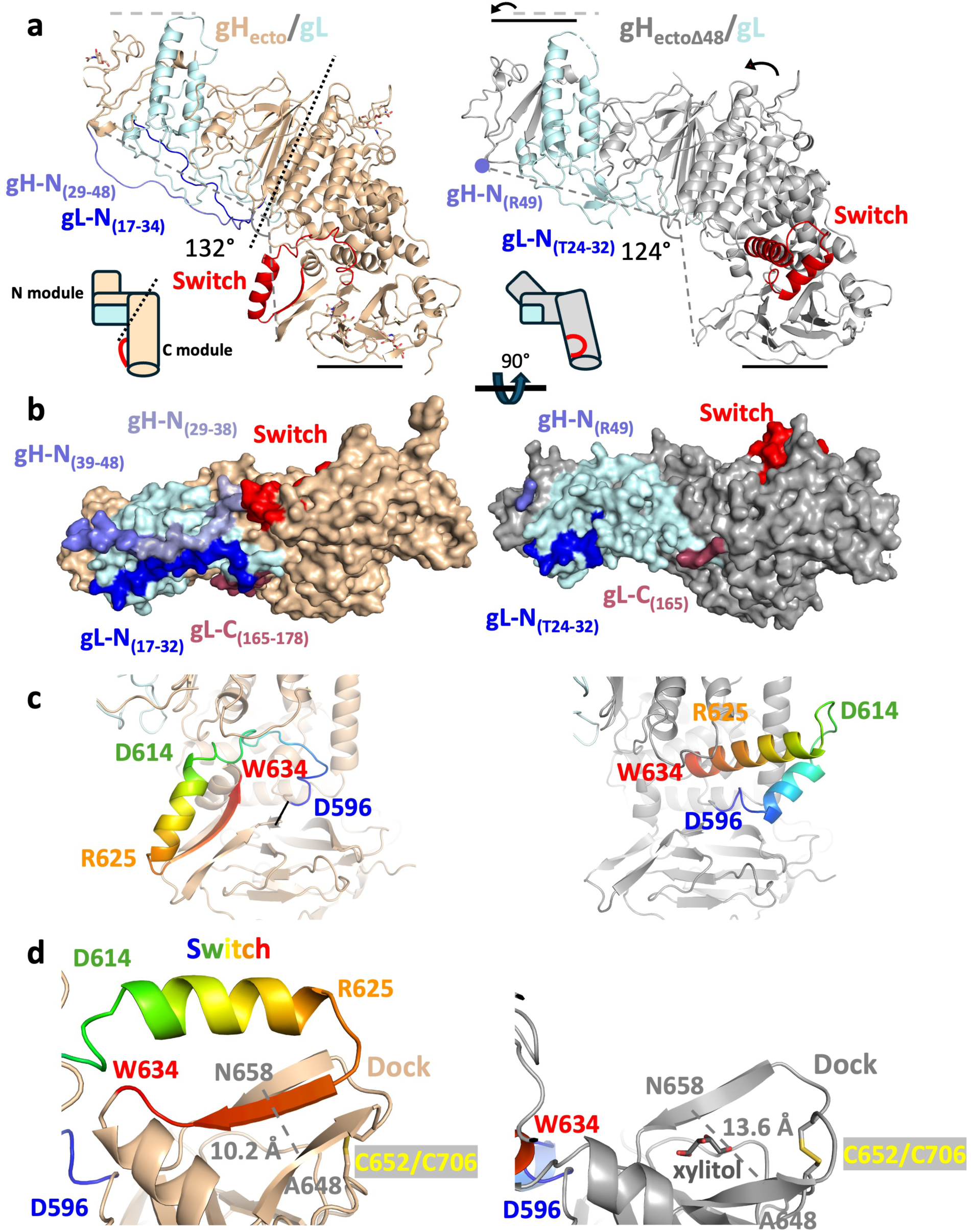
Comparison of the structures of the HSV-2 gH_ecto_/gL (pre-activation state) and the gH_ectoΔ48_/gL (activated state). **a)** Ribbon diagrams of the two structures. gH is shown in wheat and grey; gL is shown in light blue. The N termini of gH and gL resolved in the gH_ecto_/gL structure are shown in slate and blue, respectively. The switch region of gH is shown in red. The global differences in the overall conformations are shown as differences in the height and the angle. The approximate boundary between the N- and the C-terminal modules of gH/gL is shown as a dashed line. Insets depict the two states schematically. **b)** Surface diagrams, colored as in A. The N terminus of gH is colored according to the resolution of the cryoEM maps used in tracing, aa 39-48 (4.19 Å, slate) and aa 29-38 (5.56 Å, light blue). The C terminus of gL resolved in the gH_ecto_/gL structure is shown in maroon. **c)** The switch motif (residues 596-636, color ramped from blue to red) adopts a loop-strand-helix conformation in the gH_ecto_/gL structure and a helix-loop-helix conformation in the gH_ectoΔ48_/gL structure. The loop is made up of residues 599-610, the helix is composed of residues 611-626, and the strand is made up of residues 628-636. Select residues are shown in colors matching their location. **d)** In the gH_ecto_/gL structure, the β-strand of the switch is positioned between two β-strands of the dock (residues 644-659) completing a three-strand β-sheet. In the gH_ectoΔ48_/gL structure, a xylitol molecule occupies the larger space between the two β-strands of the dock. The distance between the two β-strands of the dock is shown as a dashed line. PDB ID for activated gH/gL, 3M1C.

In addition to distinct domain orientations, the gH_ecto_/gL structure has two major conformational differences from the gH_ectoΔ48_/gL structure. First, the N-terminal regions of gH and gL that were missing or unresolved in the gH_ectoΔ48_/gL structure, form an X-shaped feature covering one face of the N-terminal H1/gL module (**Figure 1B**). gH residues 29-48, missing from the gH_ectoΔ48_/gL structure, form an extended polypeptide that runs from the picket fence along the surface composed of gL loops to the tip of the gH/gL heterodimer. gH residues 39-47 were resolved in the 4.19-Å map whereas residues 29-38 were only resolved in the 5.56-Å N-terminal focus map (described below) at lower map threshold values (**Figure 1B**). gL residues 17-23, unresolved in the gH_ectoΔ48_/gL structure, also form an extended polypeptide that interacts with the N terminus of gH and with the C terminus of gL, residues 165-178, also unresolved in the gH_ectoΔ48_/gL structure (**Figure 1B**).

Second, residues 596-636, located at the C terminus of domain H2 in the arch of the boot, adopt a very different fold. In the gH_ectoΔ48_/gL structure, these residues adopt a helix-loop-helix conformation (**Figure 1C**). However, in the gH_ecto_/gL structure, they instead adopt a loop-helix-strand conformation (**Figure 1C**). The entire loop-helix-strand element is flipped ∼180 degrees relative to the gH_ectoΔ48_/gL structure, akin to a light switch (**Figure 1A, 1C, Supplemental Movie 1)**. We refer to this region as a “switch.” In the gH_ecto_/gL structure, the β-strand (residues 628-636) of the switch is docked between two β-strands, residues 644-649 and 655-659, completing a three-strand β-sheet (**Figure 1C and 1D**). We refer to this region as a “dock”. In the gH_ectoΔ48_/gL structure, the two strands are spaced farther apart, with the shortest distance Cα_A648_-Cα_N658_ of 13.2 Å (compared to 10.6 Å in the gH_ecto_/gL structure) (**Figure 1D**). A xylitol molecule, used as a crystal cryoprotectant, occupies this larger interstrand space (**Figure 1D, right**). In both structures, the dock is anchored to the H3 disc by a disulfide between residues C652 and C706 (**Figure 1D**). Mutagenesis targeting disulfide-forming cysteines in HSV-1 and HSV-2 gH/gL showed that these cysteines are important for fusion^25^.

The presence of the intact gH N terminus along with the observed structural differences suggests that the cryoEM structure of gH_ecto_/gL represents the pre-activation state of the gH/gL ectodomain. A 6.11-Å reconstruction of the unbound gH_ecto_/gL heterodimer, which resembles that of the CHL27-bound gH_ecto_/gL, further supports this notion (**Extended Data Figure 4A and B**).

### CHL27 Fab primarily interacts with the N terminus of gH

To visualize the gH_ecto_/gL-CHL27 interface, we used a focused classification and refinement of the complete particle set by imposing an N-terminal module mask (**Extended Data Figure 5**). The epitope-binding surface was resolved well enough in the 4.19 Å map to permit tracing of the N-terminal gH binding epitope (residues 38-47, slate), and N-terminal gH residues 29-37 (light blue) were placed into lower-confidence density in the 5.56 Å map (**Figures 1B, and 2A**).

Both V_H_ and V_L_ of CHL27 Fab contribute to interactions with gH and gL (**Figure 2B, 2C**). Residues 43-47 of gH fit into a vertical groove made up of the complementarity determining regions (CDRs) 1 and 3 of the V_H_ chain, CDR-H1 and CDR-H3 (**Figure 2B**). The CDR-L3 completes the binding site for residues 43-47 of gH at the junction between the variable regions. Additionally, CDR-L1 and CDR-L2 are close enough to gL residues 27-34 and, possibly, gH residues 39-42 to interact (**Figure 2C**), but these interactions are not sufficiently resolved to discern side chain orientations. The gH/CHL27 interactions observed in our structure are consistent with prior epitope mapping studies using a series of overlapping peptides, which mapped the CHL27 epitope to gH residues 37-47^21^.

**Figure 2.**
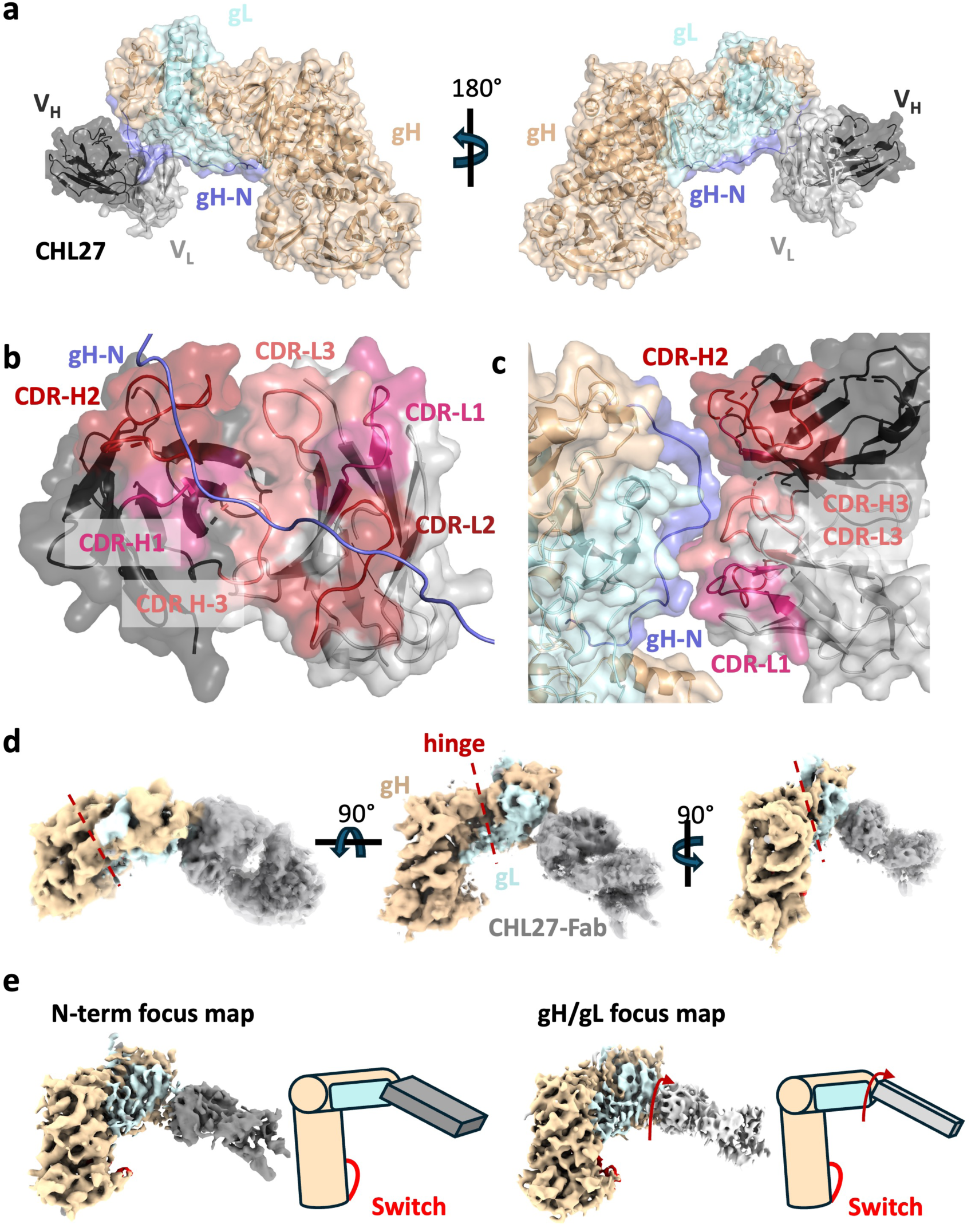
CHL27 Fab dynamically binds a membrane-distal surface of gH/gL. **a)** The CHL27 Fab light (V_L_) and heavy (V_H_) chains bind the gH/gL N terminus at residues 43-47 of gH. **b)** Complementarity-determining regions (CDRs) of both the heavy and light chains of CHL27 interact with the gH N terminus. Most contacts with gH residues 43-47 occur with V_H_. **c)** V_L_ is in proximity to gH and gL residues. CDR-L3 is proximal to gL residues 27-34, yet rigid-body fitting and refinement did not give conclusive evidence of amino-acid level interactions. **d)** CHL27-Fab is dynamically bound to the gH/gL heterodimer. **e)** Focused classification and refinement of the gH/gL-Fab complex using N-terminal or a full gH/gL mask reveals a mobile Fab that rotates and twists with respect to the gH/gL heterodimer. gH N terminus: slate, CHL27 V_L_: light gray, CHL27 V_H_: dark gray, CDR-H1: pink, CDR-H2: brick, CDR-H3: light salmon, CDR-L1: pink CDR-L2: brick, CDR-L3: light salmon.

The low resolution of CHL27 and its epitope hinted at a highly dynamic binding mode. To identify possible global conformational heterogeneity, we reconstructed the entire 820,197-particle dataset into a 5.85-Å consensus map and performed a 3D Flexibility analysis (3DFlex) in cryoSPARC^26^. This analysis generated a volume series describing the motion of CHL27 Fab and its effects on the heterodimer. As the Fab moves relative to gH/gL, the N-terminal module also flexes about a hinge located between the gL helical hairpin and the gH picket fence (**Figure 2D, Supplemental Movies 2 and 3**). Comparison of reconstructions from the two different focused classifications recapitulated the motions suggested from 3DFlex analysis (**Figure 2E**). The highly mobile nature of the gH_ecto_/gL-CHL27 interaction likely contributed to the low local resolution of the CHL27 Fab relative to gH_ecto_/gL (**Extended Data Figure 2B**).

### The gH/gL-CHL27 complex exists as an ensemble of conformational states spanning a pre-activation-to-activated continuum

To uncover conformational differences within the gH_ecto_/gL heterodimer that might have otherwise been overshadowed by the large-scale movements of the Fab, we generated a masked gH/gL-only cryoEM map and examined it for heterogeneity using 3D Variability analysis (3DVA)^27^. 3DVA resolved a series of 20 volumes representing continuous conformational heterogeneity in the gH_ecto_/gL ectodomain (**Extended Data Figure 6, Supplemental Movies 4 and 5**). Thus, the gH_ecto_/gL-CHL27 complex exists as a conformational ensemble in solution. The conformational states identified by this analysis were independently validated using an orthogonal machine-learning algorithm as implemented in CryoDRGN^28^ (**Supplemental Movie 6**). Conformational state 1 on one end of the continuum matches the 4.19-Å cryoEM reconstruction of the pre-activation state of gH_ecto_/gL (**Figure 3A, left and Extended Data Figure 6, top left**). Conversely, conformational state 20 on the opposite end of the continuum resembles the activated state captured in the crystal structure of gH_ectoΔ48_/gL (**Figure 3A, right and Extended Data Figure 6, bottom right**). Finally, one of the intermediate conformational states matches the 5.56-Å cryoEM map generated by imposing an N-terminal module mask (**Figure 3A, middle and Extended Data Figure 6, middle**).

**Figure 3.**
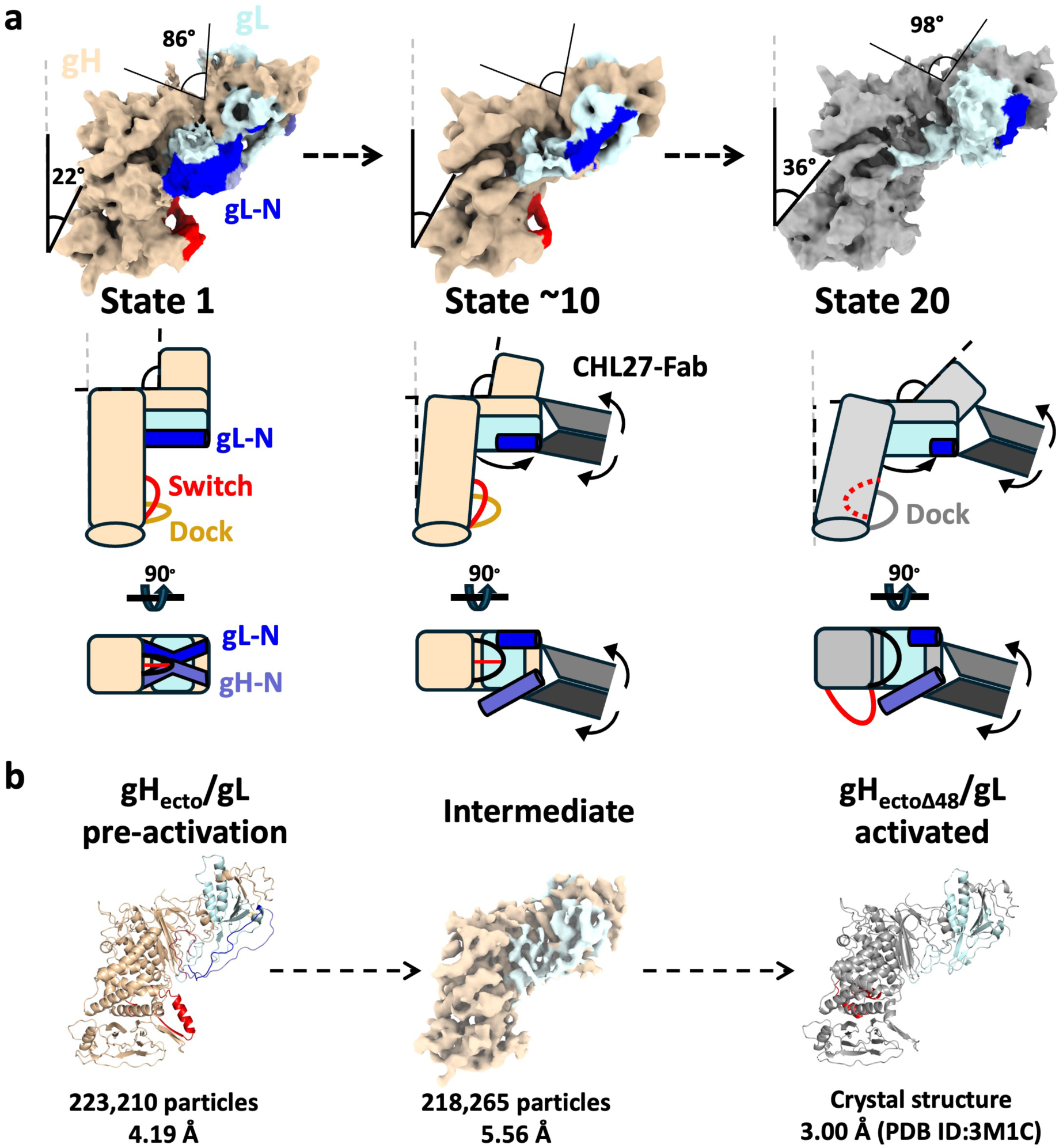
gH/gL-CHL27 complex exists in a conformational equilibrium. **a)** Local and global conformational rearrangements as revealed by 3DVA. The conformational continuum is bookended by states that correspond to pre-activation (left) and post-activation gH/gL states. Locally, N-terminal gL density translates across the N-terminal module (top asterisk) as the helical density corresponding to the switch disappears (bottom asterisk). Globally, the angle between the H2 central pillar and the vertical increases, as does the angle between the gD-binding region and the rest of the N-terminal module. Electron densities are represented diagrammatically below the continuum, as are the structures determined for gH_ecto_/gL and gH_ectoΔ48_/gL. Model for the sequential activation of gH/gL by CHL27: (*Left*) gH/gL in its pre-activation state stands tall, with gH and gL N termini making contact with the picket fence. (*Center*) As the Fab binds and applies force, gH and gL disengage from their binding surfaces. Angles from the vertical and between the N- and C-terminal modules subsequently increase, leading to a “tilt” as seen in the gH_ectoΔ48_/gL structure. (*Right*) The distance between the two strands of the dock increases. Density corresponding to the switch motif disappears from below the N-terminal module, indicating its refolding and repositioning into the helix-loop-helix configuration captured by the gH_ectoΔ48_/gL crystal structure. **b)** Structures solved for the pre-activation and activated states of the gH/gL ectodomain. PDB ID: pre-activation gH_ecto_/gL (9CZG), activated gH_ectoΔ48_/gL (3M1C).

Both local and global conformational rearrangements occur during the transition from state 1 to state 20 along the continuum (**Figure 3B and Extended Data Figure 6**). Locally, the cryoEM density corresponding to the gL N terminus, residues 17-30, shifts across the N-terminal module. Concurrently, the density corresponding to the switch, residues 596-636, disappears from its position below the N-terminal module (**Figure 3B**, **Supplemental movies 4 and 5**). Globally, the angle between the gD-binding region and the rest of the N-terminal module increases from 86° to 98° (**Figure 3**). The entire gH/gL heterodimer tilts away from the vertical, evident from the angle between the sole of the gH/gL boot and the vertical increasing from 22° to 36°.

The resemblance between the terminal states 1 and 20 and the structures of the pre-activation and activated states of gH/gL, respectively, suggests that the observed continuum captures conformational transitions in gH/gL as it refolds from the pre-activation to activated state. The presence of the activated state, previously captured in crystals, in our cryoEM data further supports this notion.

## DISCUSSION

gH/gL is a conserved component of the herpesviral entry machinery that transmits the receptor engagement signal from gD to gB thereby coupling host-receptor binding to membrane fusion during host entry. To achieve this, it must undergo conformational changes that somehow activate the fusogen, gB. The nature of these conformational changes has remained elusive, however. Here, we determined the structure of the gH_ecto_/gL complex in its pre-activation state, prior to interactions with the receptor-bound gD. To achieve this, we preserved the N terminus of gH and used a neutralizing Fab CHL27 to stabilize its conformation and enable particle alignment, a strategy that previously succeeded with other macromolecules^29^. The other known conformational state of gH/gL, captured in the crystal structure of the gH_ectoΔ48_/gL construct lacking the gH N terminus^19^, has been proposed to represent the activated state of the heterodimer^23^. Indeed, the match between this structure and one of the terminal states in the conformational ensemble observed in the cryoEM data supports this idea.

### The N termini of gH and gL stabilize the pre-activation state

Our cryo-EM reconstruction of the gH_ecto_/gL resolved the N termini of gH and gL that were either missing or disordered, respectively, in the gH_ectoΔ48_/gL crystal structure^19^. In the gH_ecto_/gL structure, these segments interact to form an X-shaped feature covering one face of the N-terminal module (**Figure 1B**). The structure clarifies why antibodies CΔ48L3 and CHL18, which bind epitopes within the C terminus of gL (173 to 183 and 208 to 219, respectively), interact with gH_Δ48_/gL variants but not with intact gH/gL^21,23,24^. This was puzzling because the C terminus of gL, which was unresolved in the gH_ectoΔ48_/gL crystal structure, was expected to be surface exposed. The gH_ecto_/gL structure shows that the C terminus of gL is covered by the N terminus of gL, which interacts with the N terminus of gH (**Figure 1B**). Thus, the epitopes of CΔ48L3 and CHL18 should be hidden in intact gH/gL, which also explains why these antibodies are non-neutralizing^21,23,24^. The lack of the N terminus of gH in the gH_ectoΔ48_/gL destabilizes interactions between the N and C termini of gL, exposing these epitopes. Indeed, despite being non-neutralizing, CΔ48L3 and CHL18 block membrane fusion in the presence of gH_Δ48_/gL^21,23,24^.

Our analysis suggests that the N termini of gH and gL both help stabilize gH/gL in its high-energy pre-activation state (**Figure 1B**). This explains why the gH_Δ48_/gL variant lacking the gH N terminus can support fusion in a cell-cell fusion assay when co-transfected with gB, be it in the presence or absence of gD^23^.In the absence of the gH N terminus, the pre-activation state of gH_ectoΔ48_/gL is stabilized by the N terminus of gL and can be activated by gD similarly to the intact gH/gL. However, lacking the stabilizing interactions provided by the gH N terminus, the pre-activation state of gH_Δ48_/gL is less stable and can be triggered even in the absence of gD, albeit less efficiently.

### The gD-binding site in gH/gL and the role of CHL27

Activation of gH/gL requires binding to a receptor-bound gD. A panel of antibodies, including CHL27, that bind the N terminus of gH block gD-gH/gL binding. Thus, the gD-binding site in gH/gL at least partially overlaps the epitope of CHL27, residues 43-47 within the N terminus of gH. A more precise identification of the gD-binding site awaits the structure of the gD/receptor-gH/gL complex. Such a structure would also clarify the triggering of conformational rearrangements in gH/gL.

Initially, we hypothesized that binding of CHL27 to the N terminus of gH would help stabilize the pre-activation state of gH/gL. Serendipitously, this complex yielded an ensemble of conformational states spanning the range from pre-activation to intermediate and activated. Partial stabilization of the pre-activation state by CHL27 could be due to experimental conditions – brief crosslinking at room temperature after a short incubation of the two components of the complex on ice – that failed to trap all gH/gL in pre-activation state. However, we cannot yet rule out the possibility that CHL27, instead, triggered gH/gL to refold from its pre-activation to activated state, mimicking receptor-bound gD. In this latter case, neutralization by CHL27 could be due to non-productive conversion of gH/gL to its activated form in the absence of receptor. Regardless, the gH/gL-CHL27 complex allowed us to capture and characterize the conformational continuum of gH/gL states that lead to activation of gB.

### Flipping of the switch in gH activates gB

The most striking difference between the gH_ecto_/gL and the gH_ectoΔ48_/gL structures, which represent the pre-activation and activated states, respectively, is the refolding and flipping of the switch element in domain H2. The switch adopts a loop-helix-strand conformation in the pre-activation state but refolds into a helix-loop-helix conformation in the activated state (**Figure 1C, 1D**). This refolding is accompanied by a ∼180-degree flipping.

What is the role of switch? In the gH_ecto_/gL structure, it is docked in the middle of a β-sheet and tucked under the N-terminal module. By contrast, in the gH_ectoΔ48_/gL structure, the switch protrudes out from the wide surface near the putative epitope of the antibody LP11 that blocks gH/gL-gB interactions^19^. We hypothesize that in its flipped conformation, the switch would preclude gH/gL-gB interactions. This raises the question of how gH/gL and gB interact at different stages of fusion.

### gH/gL-gB interactions

Our prior work using split luciferase reporters showed that gH/gL and gB interact before and during fusion^22^. We hypothesize that in its pre-activation state, gH/gL ectodomain interacts with the gB ectodomain and that this interaction helps stabilize the metastable prefusion form of gB. Conversely, interactions with the prefusion gB could help stabilize the pre-activation state gH/gL, locking both glycoproteins in a mutually inhibited complex on the surface of a virion or an infected cell. We also showed that in the absence of gL, not only all gH but also a large portion of gB is retained in the ER, suggesting that gH/gL and gB are likely transported as a complex to the cell surface^22^. Flipping of the switch during transition of gH/gL to its activated state would disrupt interactions with the prefusion gB ectodomain, providing a mechanism for releasing gB and allowing it to refold into the postfusion conformation. This model implies that rather than being activated by interactions with gH/gL, gB is released from an autoinhibited complex with gH/gL. In support of this idea, complexes of prefusion gB bound to gH/gL have been observed on the surface of HCMV virions by cryoelectron tomography and immunoprecipitation^30,31^. Thus, we posit that gH/gL serves as a “lynchpin,” constraining gB in its metastable prefusion state prior to host-receptor binding by gD.

How do gH/gL and gB maintain interactions during fusion, i.e., post-triggering? Previously, we found that the intraviral (cytoplasmic) regions of gB and gH regulate the membrane fusion process and proposed the “clamp-and-wedge” model^32^. The cytoplasmic domain (CTD) of gB forms a membrane-interacting inhibitory clamp that helps restrain the gB ectodomain in its prefusion conformation, presumably, through interactions with the transmembrane and membrane-proximal regions on the opposing side of the membrane^32^. The cytoplasmic tail (CT) of gH disrupts the inhibitory gB CTD clamp by acting as a wedge, which helps trigger the fusogenic refolding of the gB ectodomain. Using mutagenesis and modeling, we narrowed down the interacting regions to a hydrophobic pocket at the interprotomeric interface in the gB CTD trimer and residue V831 of gH CT^33^.

We hypothesize that the loss of interactions between the ectodomains of gH/gL and gB coincides with the disruption of the gB CTD clamp by the insertion of the gH CT wedge **(Figure 4).** This model helps explain how gB and gH/gL can maintain interactions before and during fusion despite undergoing conformational changes. We propose that prior to fusion, gB and gH/gL primarily interact through their ectodomains whereas during fusion, the interactions are primarily mediated by the cytoplasmic domains. Thus, the activation process involves coordinated actions across both the ectodomains and the cytoplasmic domains of gH/gL and gB. Future studies will unveil the precise order of these events and their coordination. Indirect evidence of gB-gH/gL complexes before and during fusion would be greatly bolstered by determining the structure of that complex.

**Figure 4.**
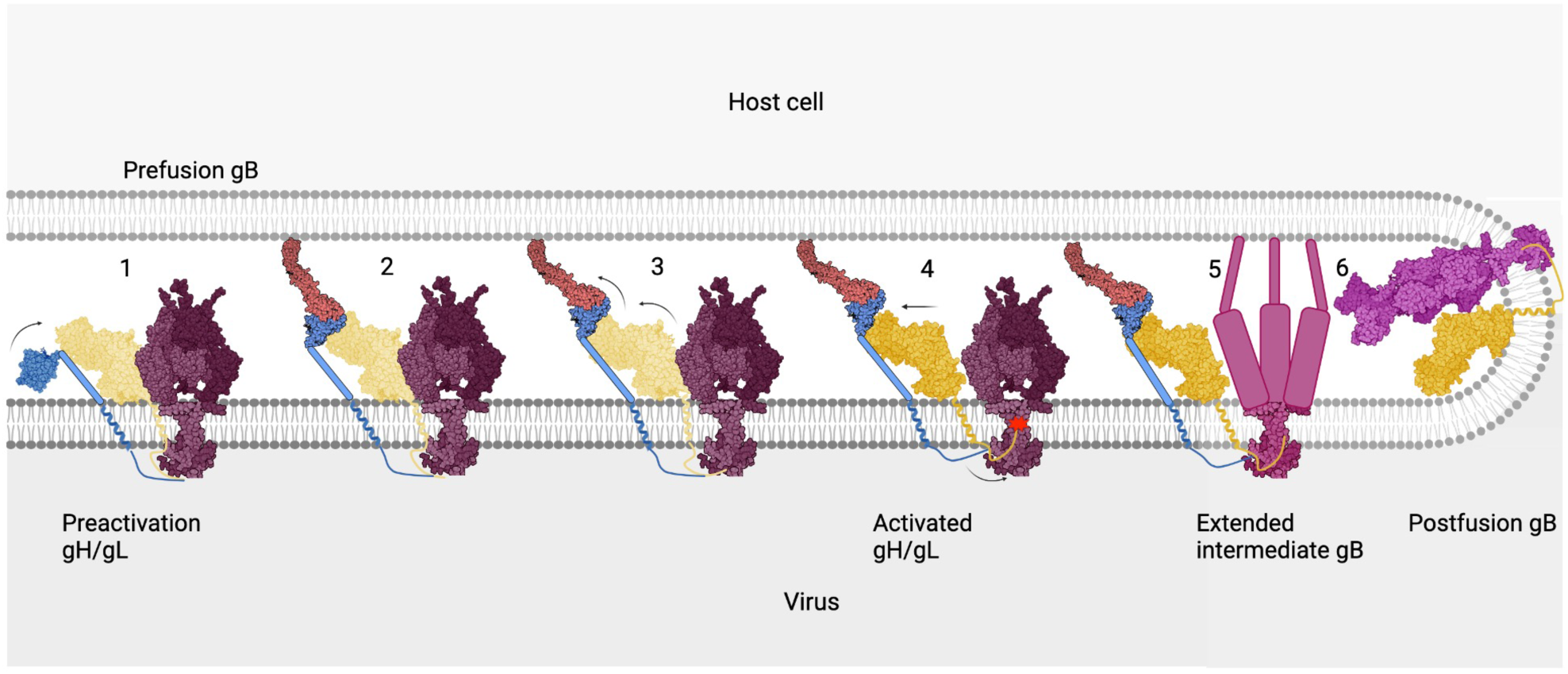
Stepwise model for gH/gL-mediated activation of gB. 1) Pre-activation gH/gL interacts with prefusion gB through their ectodomains. 2) Binding to receptor causes gD to bind the N terminus of gH/gL. 3) Binding of the gD/receptor complex to the N terminus of gH/gL causes a series of conformational changes in gH/gL that propagate throughout the heterodimer, culminating in the release and refolding of the switch. 4) In its activated state, the ectodomain of gH/gL no longer interacts with the prefusion gB. On the opposing side of the membrane, the cytoplasmic tail of gH engages the cytoplasmic domain of gB. This interaction destabilizes the cytoplasmic “clamp” that releases its grip onto the ectodomain. 5) Prefusion gB refolds into its extended intermediate conformation, exposing the fusion loops that interact with the membrane of the host cell. 6) As gB continues to refold into its final postfusion conformation, the fusion loops and the transmembrane spans are brought into proximity. This action brings apposing membranes close enough to merge, completing the fusion process. gD (blue), nectin-1 (brick), pre-activation gH/gL (pale yellow), prefusion gB (bordeaux), activated gH/gL (goldenrod), intermediate gB (dark rose), and postfusion gB (magenta). PDB IDs: gD (2C36), nectin-1 (3U83), activated gH/gL (3M1C), prefusion gB (6Z9M and 5V2S), and postfusion gB (5V2S).

### Flip-switch transmission of the receptor-binding signal from gD to gB through gH/gL

We hypothesize that the continuum of conformational states observed by cryoEM here recapitulates the steps that transmit the activating signal from receptor-bound gD to prefusion gB (**Extended Data Figure 7**). We envision that gH/gL folds into a high-energy, pre-activation state stabilized by the interactions between the gH and gL N termini and, potentially, prefusion gB (**Extended Data Figure 7, step 1**). When receptor-bound gD binds to the N terminus of gH, this releases it from its contacts with the N terminus of gL and the picket fence (**Extended Data Figure 7 step 2, Supplemental Movies 4 and 5**). This action also releases the N terminus of gL, which we observe as a large shift in cryoEM density toward the top of the heterodimer (**Figures 1B and 3, Supplemental Movies 4 and 5**). The release of the N termini coincides with the tilting of domain H2 relative to domain H3 (**Figures 1C and 3**). The switch is then released by weakened inter-strand interactions when the distance between the two outer strands of the dock (residues 644-662) is widened by ∼3 angstroms, allowing it to flip and refold into the helix-loop-helix conformation and captured in the gH_ectoΔ48_/gL structure (**Figure 1C, D, Extended Data Figure 7, step 3, and Supplemental Movie 1**). In this manner, the conformational rearrangements described above propagate across the gH/gL heterodimer from the N terminus of domain H1 to the C terminus of domain H2 (**Figure 3, Supplemental Movie 1**). In this conformation, the switch juts out from the gB-binding surface of gH/gL.

### The “lynchpin” model for fusion regulation by gH/gL

These structural studies lead us to propose a lynchpin model for how the gH/gL ectodomain regulates gB fusion activity (**Figure 4**). Prior to gD binding of a host receptor, pre-activation gH/gL and prefusion gB interact through their ectodomains, helping stabilize each other (step 1). Binding to receptor causes gD to bind the N-terminus of gH/gL (step 2). Binding of gD/receptor complex to gH/gL initiates a series of conformational changes that propagate throughout the heterodimer, culminating in the release, refolding, and flipping of the switch (step 3). Once flipped, the switch disrupts the interactions between the ectodomains of gH/gL and gB. In its activated state, the ectodomain of gH/gL no longer binds the prefusion gB ectodomain. Meanwhile, on the opposing side of the membrane, the gH CT wedges into the gB CTD, which destabilizes the clamp and causes it to release its grip on the gB ectodomain (step 4). The loss of the stabilizing interactions with the gH/gL ectodomain allows the prefusion gB to refold into the extended intermediate (step 5) and, ultimately, (6) to its stable, low-energy postfusion conformation, completing the fusion process. We hypothesize that refolding of the gH switch from loop-helix-strand to helix-loop-helix is thermodynamically favorable. Thus, gH/gL is a “one-shot” machine that irreversibly refolds from pre-activation to activated form, similarly to gB, which irreversibly refolds from prefusion to the postfusion form^18^.

There are several implications to our model that potentially apply across all *Herpesviridae* subfamilies. First, the switch element, in its activated helix-loop-helix conformation, is found across all known structures of gH/gL homologs (**Extended Data Figure 8**). Therefore, we hypothesize that this element has a conserved role in the gB activation mechanism by gH/gL. Thus, we suspect that our proposed model for how gH/gL regulates the activity of gB could be generally applicable across the entire family *Herpesviridae*. Finally, the identification of the switch mechanism for gB activation by gH/gL points to a target for subunit-based vaccines and treatment efforts, possibly applicable to all herpesviral infections.

## MATERIALS AND METHODS

### Expression and purification of the recombinant HSV-2 gH/gL

Recombinant baculovirus (pTC605) encoding HSV-2 gH ectodomain, residues 1-803, with a C-terminal His_6_ tag and full-length HSV-2 gL, residues 1-224, was generated using the pFastBac Dual system as previously described (**Extended Data Figure 8A**)^34^. Virus titer was increased by two rounds of serial passaging in Sf9 cells to passage 3 (P3). For protein expression, 4.5-6-L cultures of Sf9 cells were infected with 8 mL of virus per 1 L of cells at a density of 2×10^6^ cells/mL. Supernatants containing the secreted gH_ecto_/gL were collected 72-80 hours post-infection and centrifuged at 4,000 g for 40 minutes in a Sorvall Lynx 4000 floor centrifuge (ThermoFisher) using a swinging bucket rotor. The supernatant was decanted and filtered using 0.2 μm polyethylenesulfonate (PES) bottle top vacuum filters. 100 mM phenylmethylsulfonylfluoride (PMSF) was added to the supernatant at a 1:1000 dilution for a working concentration of 100 μM.

The supernatant was concentrated from 4.5 or 6 L to 1.2 L using a tangential flow filtration (TFF) system as previously described^35^. Supernatant was incubated with 3-4 mL Ni-NTA Excel resin (GE/Cytiva) pre-equilibrated in phosphate buffered saline (PBS) for 45 minutes at 4 °C in 50-mL conical tubes under nutation. The resin was pelleted by centrifugation at 1,000 g for 5 minutes in a tabletop centrifuge using a swinging bucket rotor, and the supernatant containing the flow-through fraction was carefully removed. The resin was washed using gravity flow with 60-80 mL wash buffer 1 (25 mM Tris, pH 7.9, 150 mM NaCl, 10 mM imidazole); 60-80 mL wash buffer 2 (25 mM Tris, pH 7.9, 500 mM NaCl, 20 mM imidazole); and 60-80 mL of wash buffer 3 (25 mM Tris, pH 7.9, 150 mM NaCl, 20 mM imidazole) prior to elution with 25 mL elution buffer (25 mM Tris, pH 7.9, 150 mM NaCl, 140 mM imidazole) into a 50-mL conical tube containing 50 μL of 0.5 M ethylenediaminetetraacetic acid (EDTA) for a final concentration of 1 mM. The eluate fraction was concentrated using 100 kDa MWCO 15 mL spin concentrators (Amicon/Millipore) to 500 μL, centrifuged at 21,000 g at 4 °C for 10 minutes to remove aggregates, and injected onto a 10/300 Superose 6 column (GE/Cytiva) pre-equilibrated with size exclusion/storage buffer (25 mM HEPES, pH 7.4, 150 mM NaCl) using a Biorad NGC system. Chromatographic peaks were analyzed by SDS-PAGE (**Extended Data Figure 1B**). Relevant fractions were pooled and concentrated to ∼5 mg/mL in storage buffer with a 100 kDa MWCO 4 mL spin concentrator (Amicon/Millipore) for immediate use or snap-frozen in 25-μL aliquots for future use.

### Purification of CHL27 Fab

CHL27 Fab was isolated using a Pierce Fab Preparation Kit (Thermo Scientific), according to the manufacturer’s protocol. Briefly, 5 mgs of murine IgG were incubated with 250 µl papain immobilized on agarose resin for 4h at 37°C. After digestion, Fab was purified using a Protein A spin column. Protein concentration was determined by absorbance at 280nm using an estimated extinction coefficient of 1.4.

### CHL27 isotyping and hybridoma sequencing

Mouse CHL27 Mab was isotyped using Pierce Rapid Isotyping Kit following the manufacturer’s protocol. Hybridoma sequencing were performed by Genscript (Piscataway, NJ). Total RNA was isolated from 2×10^6^ cells and reverse-transcribed into cDNA using isotype-specific (mouse IgG1 heavy chain, kappa light chain) anti-sense primers or universal primers following PrimeScriptTM 1st Strand cDNA Synthesis Kit’s manual. Antibody fragments of V_H_, V_L_, C_H_ and C_L_ were amplified according to the standard operating procedure of rapid amplification of cDNA ends of GenScript. The sequences of different clones were aligned, and the consensus sequence was provided (**Extended Data Figure 8B**).

### Cryo-EM sample preparation and data collection

gH_ecto_/gL/CHL27 Fab complex formation was verified by size exclusion chromatography (**Extended Data Figure 1B**). Complexes were assembled and crosslinked *in situ* during sample preparation. gH_ecto_/gL was diluted to 1.2 mg/mL using 1.25x molar excess of CHL27 Fab in 30μL of storage buffer. After 30 minutes of incubation on ice, electron microscopy-grade 25% glutaraldehyde (Invitrogen) was added to a final v/v percentage of 0.025% and incubated for 5 or 10 minutes. The crosslinking reaction was quenched with 4X quenching buffer (36 mM Tris, pH 8.0 with 0.4% w/v 2-octylpyranoglucoside (2-OG) (Affymetrix)) for a final Tris and detergent concentration of 9 mM and 0.1%, respectively. Sample was immediately applied to glow-discharged holey C-flat grids (1.2/1.3, 4C 400 mesh Cu grids Mitegen Cat #:M-C413) and vitrified in nitrogen-cooled liquid ethane with a Gatan CP3 cryoplunger using blot setting 4.

Samples were screened on a Talos Arctica microscope (ThermoFisher) with a K3 camera (Gatan) operated at 200 kV at the Harvard Cryo-EM center and the highest quality grids were imaged on a Titan Krios Microscope (ThermoFisher) with a K3 camera (Gatan) at the New York Structural Biology Center (NYSBC) National Center for CryoEM Access and Training (NCCAT) operating at 300 kV acceleration voltage. 50-60 frame movies were collected in counting mode at 105,000x magnification with a total dose of ∼65 e^-^/Å^2^ and a pixel size of 0.825 Å. In total 29,989 movies were collected with a defocus range of -1.2 - -2.2 μm. Movies were motion-corrected and dose-weighted via the NCCAT automated pipeline using MotionCor2 ^36^, and contrast transfer function (CTF) parameters were estimated using CTFFIND4 ^37^ (representative micrograph, **Extended Data Figure 9A**). Some of this work was performed at the National Center for CryoEM Access and Training (NCCAT) and the Simons Electron Microscopy Center located at the New York Structural Biology Center, supported by the NIH Common Fund Transformative High Resolution Cryo-Electron Microscopy program (U24 GM129539,) and by grants from the Simons Foundation (SF349247) and NY State Assembly.

### Image processing and 3D reconstructions

Data processing and analysis was performed chiefly using the cryoSPARC^38^ suite. Templates were generated from prior low-resolution density maps from images collected at the Harvard CryoEM Center (11,286 50-frame movies were collected on a Titan Krios at 300kV acceleration voltage in counting mode at 105,000x magnification with a total dose of ∼75 e^-^/Å^2^, a pixel size of 0.825 Å, and a defocus range of -1.2- -2.2 μm). These data were processed using RELION’s^39^ processing pipeline. The templates generated by this initial analysis were used to pick 6,862,651 total particles (**Extended Data Figure 5**). After successive 2D classification analyses, an initial 3D model was generated using *ab-initio* reconstruction from 73,725 particles and was further improved by heterogeneous refinement. After further 2D classification of the full particle set (representative 2D classes, **Extended Data Figure 9B**), these initial models along with “junk” 3D density maps were used for heterogeneous refinement. A subsequent heterogeneous refinement step combined two 3D classes that corresponded to two distinct “views” of the gH_ecto_/gL/CHL27 complex. The density map corresponding to the full complex (820,197 particles) was refined using homogenous refinement followed by nonuniform refinement. This density map (5.85 Å resolution) was used for cryoSPARC 3D Flexibility and 3D Variability analyses^26,27^ as well as cryoDRGN continuous heterogeneity analysis^28^. This map was then further improved by focused 3D classification using two different masks: an N-terminal-module-focused mask and a full gH_ecto_/gL mask. Although the N-terminally focused 3D classification only marginally improved the overall resolution to 5.56 Å (GFSC curves, **Extended Data Figure 9C**) (218,265 particles), it enabled tracing of the gH and gL N termini. The gH_ecto_/gL-focused 3D classification significantly improved the resolution to 4.33 Å for the gH_ecto_/gL/CHL27 complex (223,210 particles). The resolution was further improved to 4.19 Å by masking away the Fab density and using local refinement of only gH_ecto_/gL. Reference-based motion correction was attempted on these final electron density maps but did not result in a meaningful improvement of the density map quality. Data collection and refinement statistics are reported in **Extended Data Table 1**.

### Model building

The crystal structure of the N-terminally truncated gH/gL (gH_ectoΔ48_/gL, PDBID:3M1C) was used for initial model building. It was fit into the electron density in ChimeraX^40^, and the model was manually refined using COOT^41^ and ChimeraX ISOLDE module^42^. Models were real-space refined in PHENIX^43^ against the gH_ecto_/gL-focused and refined map (4.33 Å, 223,210 particles) and against a combination of the N-terminally focused map (5.56 Å, 218,265 particles) and the gH_ecto_/gL-only locally refined map (4.19 Å, 223,210 particles) using PHENIX’s combine_focused_maps module^43^. A homology model of CHL27 Fab was generated using the ROSIE2 server^44^ and fit into the N-terminally focused map (5.56 Å, 218,265 particles) in ChimeraX. The homology model of the CHL27 F_v_ region was further rigid-body fitted into the density in COOT. The Fab was then rigid-body refined against the combined map. Once the gH_ecto_/gL and the CHL27 Fab models were merged into a single model, real-space refinement of the residues at the gH_ecto_/gL/CHL27 interface was done to prevent clashes using COOT and ISOLDE. The final model was refined against the combined focused map using real-space refinement to generate the statistics report (**Extended Data Table 1**).

## Supporting information

Legends for Supplemental Movies

## Acknowledgments

We thank Carmen Rivera for help with protein purification experiments. We thank Patricia Spear (Northwestern University) for the gift of plasmids. The content is solely the responsibility of the authors and does not necessarily represent the official views of the National Institutes of Health. We dedicate this publication to the memory of Roselyn J. Eisenberg (University of Pennsylvania).

## Author contributions

Experimental design: G.L.G.-D.P. and E.E.H.

Experiment execution: G.L.G.-D.P. and R.M.W.

Generation and characterization of reagents: D.A., T.M.C., W.T.S., and G.H.C.

Supervision of project: E.E.H.

Funding acquisition: E.E.H

Data collection and processing: G.L.G.-D.P. and R.M.W.

Data analysis, hypothesis generation, model generation: G.L.G.-D.P. and E.E.H.

Manuscript-writing (first draft); G.L.G.-D.P.

Manuscript-writing (subsequent drafts): G.L.G.-D.P and E.E.H.

Manuscript edits and finalization: All authors.

## Data and materials availability

Structure model and maps have been deposited to EMDB using the following accession numbers: 9CZG – gH/gL only, 9CZR – gH/gL-CHL27 complex.

## Supplementary Materials

Movies S1 to S6

---

**Extended Data Figure 1.**
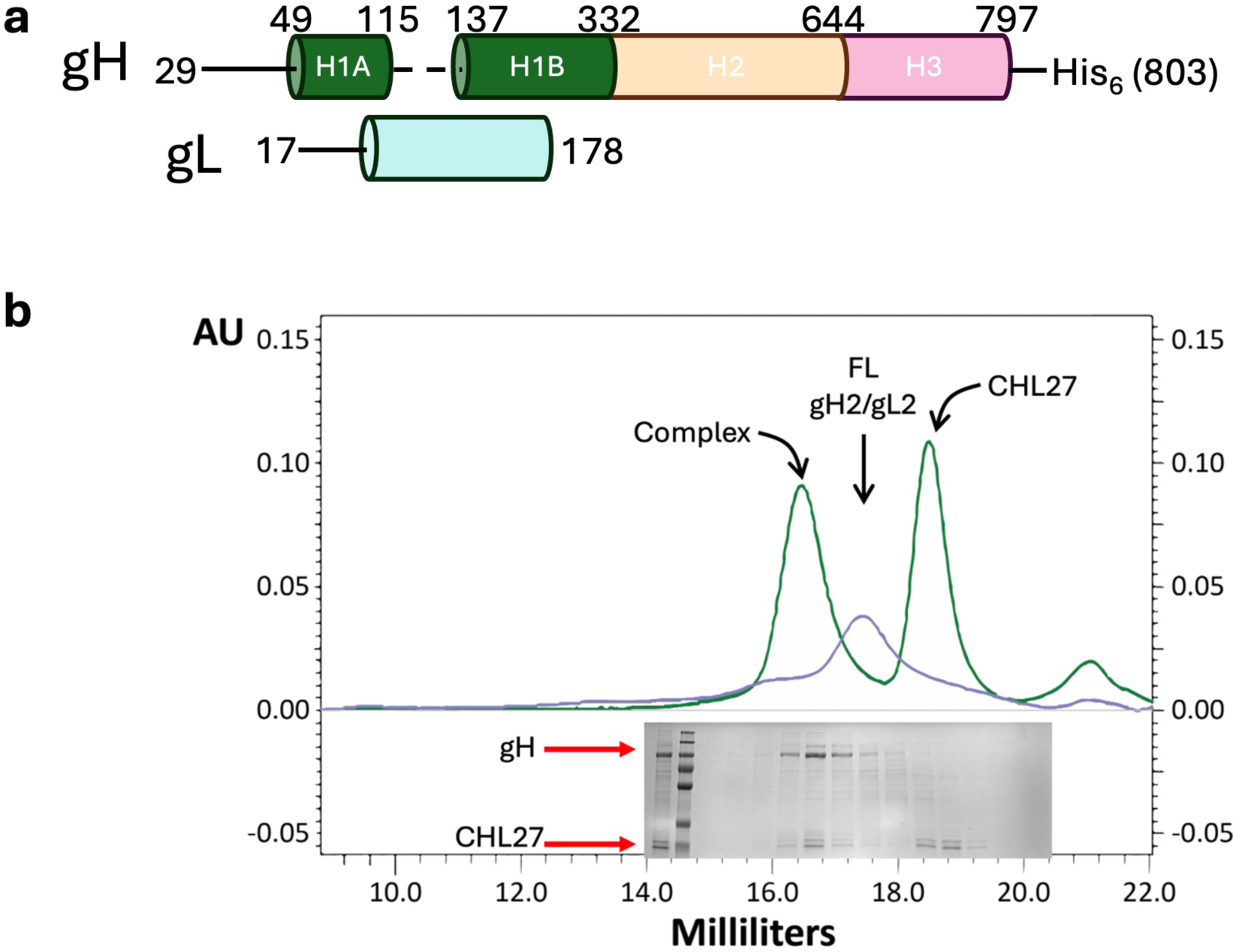
Domain architecture and purification of gH_ecto_/gL-CHL27 complex. **a)** Domain architecture of gH/gL_FL_. Domain diagram shown here represents density visible in our EM reconstruction; dotted line indicates missing density. **b)** Chromatographic and SDS-PAGE analysis confirms that gH_ecto_/gL and CHL27_Fab_ form a complex. Green trace, complex; slate trace, gH_ecto_/gL alone. gH_ecto_/gL structure resembles large-scale features of the gH_ectoΔ48_/gL structure.

**Extended Data Figure 2.**
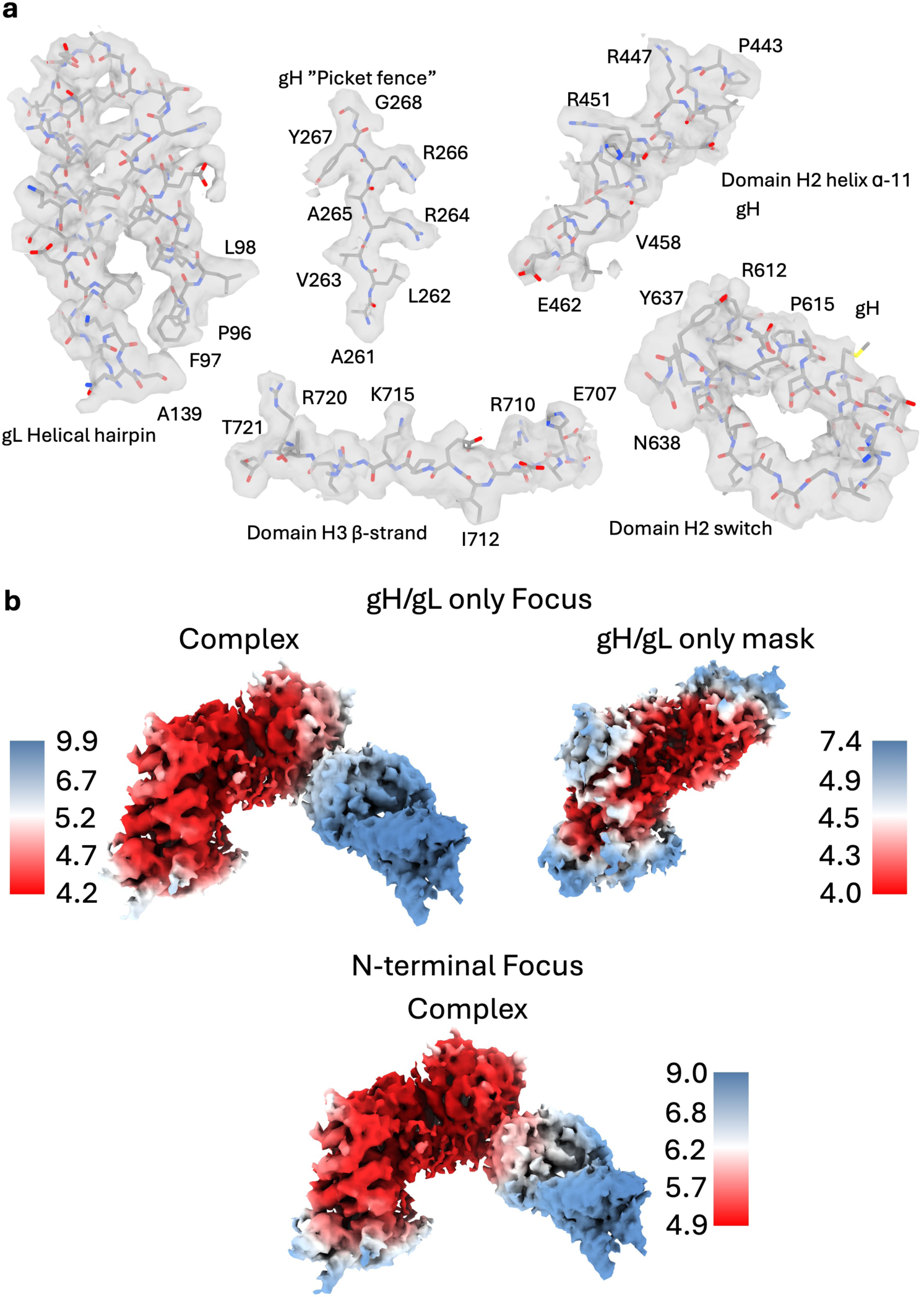
EM density quality and local resolution of gH_ecto_/gL-CHL27 complex. **a)** Density quality across gH/gL was high enough to confidently place side chains despite the relatively low overall resolution of 4.19Å. All volumes were contoured in ChimeraX to 0.86 except for the switch motif, which was contoured to 0.38 to visualize the mobile helix. **b)** Local resolution of full complex and masked gH/gL-only maps. Top: gH/gL focused maps. Left map is the full complex, right map generated by masking the Fab and performing local refinement after particle subtraction. Bottom: N-terminal module-focused map of the complex.

**Extended Data Figure 3.**
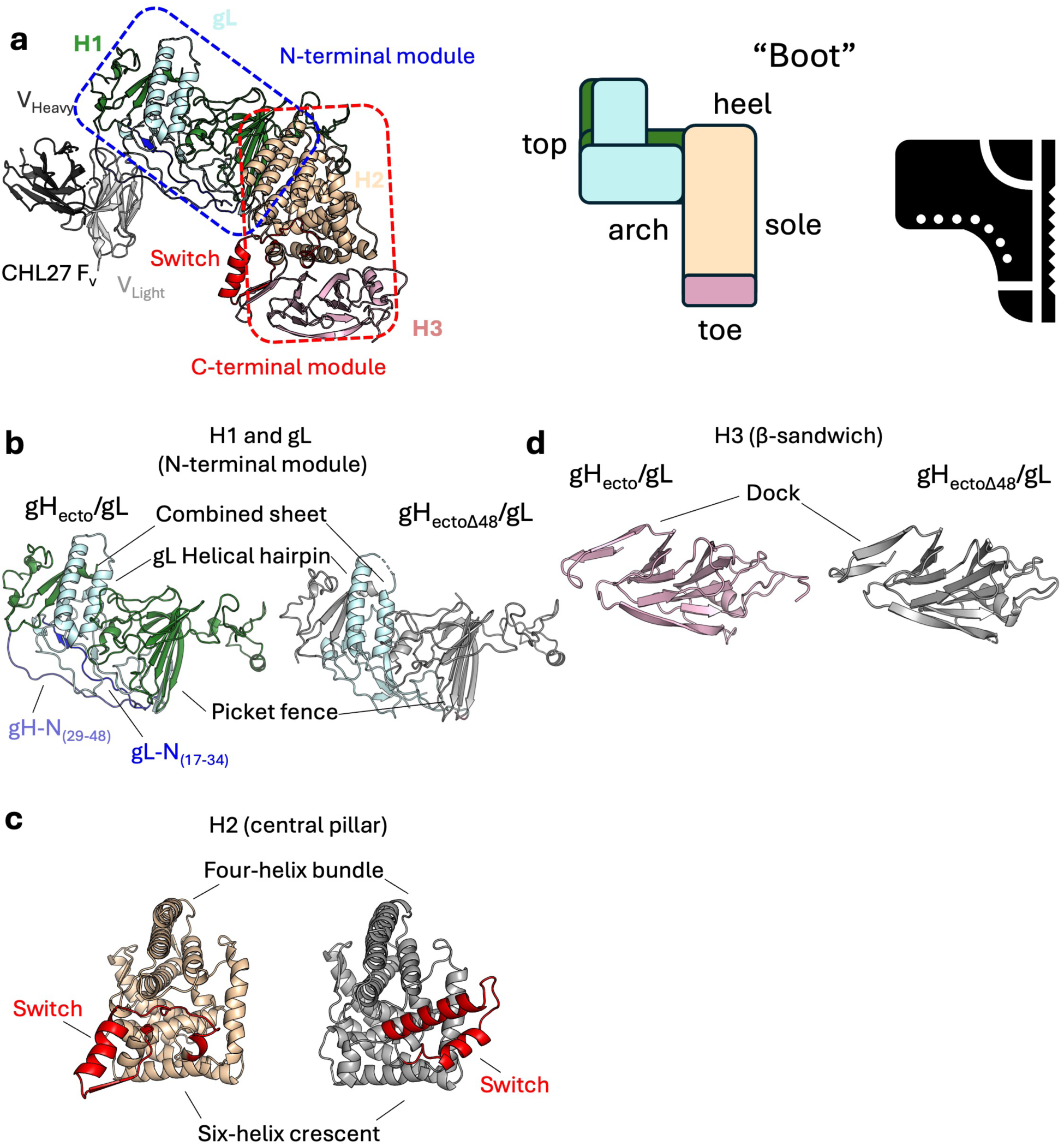
Overall structure of a gH_ecto_/gL-CHL27 Fab complex. **a)** gH_ecto_/gL structure resembles large-scale features of the gH_ectoΔ48_/gL structure, retaining its boot-like appearance. **b)** The N-terminal module contains a gH-gL combined β-sheet, gL helical hairpin, and a five-strand β-sheet picket fence. **c)** The H2 central pillar is composed of a four-helix bundle packed against a six-helix crescent. The C-terminal switch is in both a different configuration and position. **d)** Domain H3 is made up of a β-sandwich lined by loops and a helix. The switch locks into the dock in the gH_ecto_/gL structure.

**Extended Data Figure 4.**
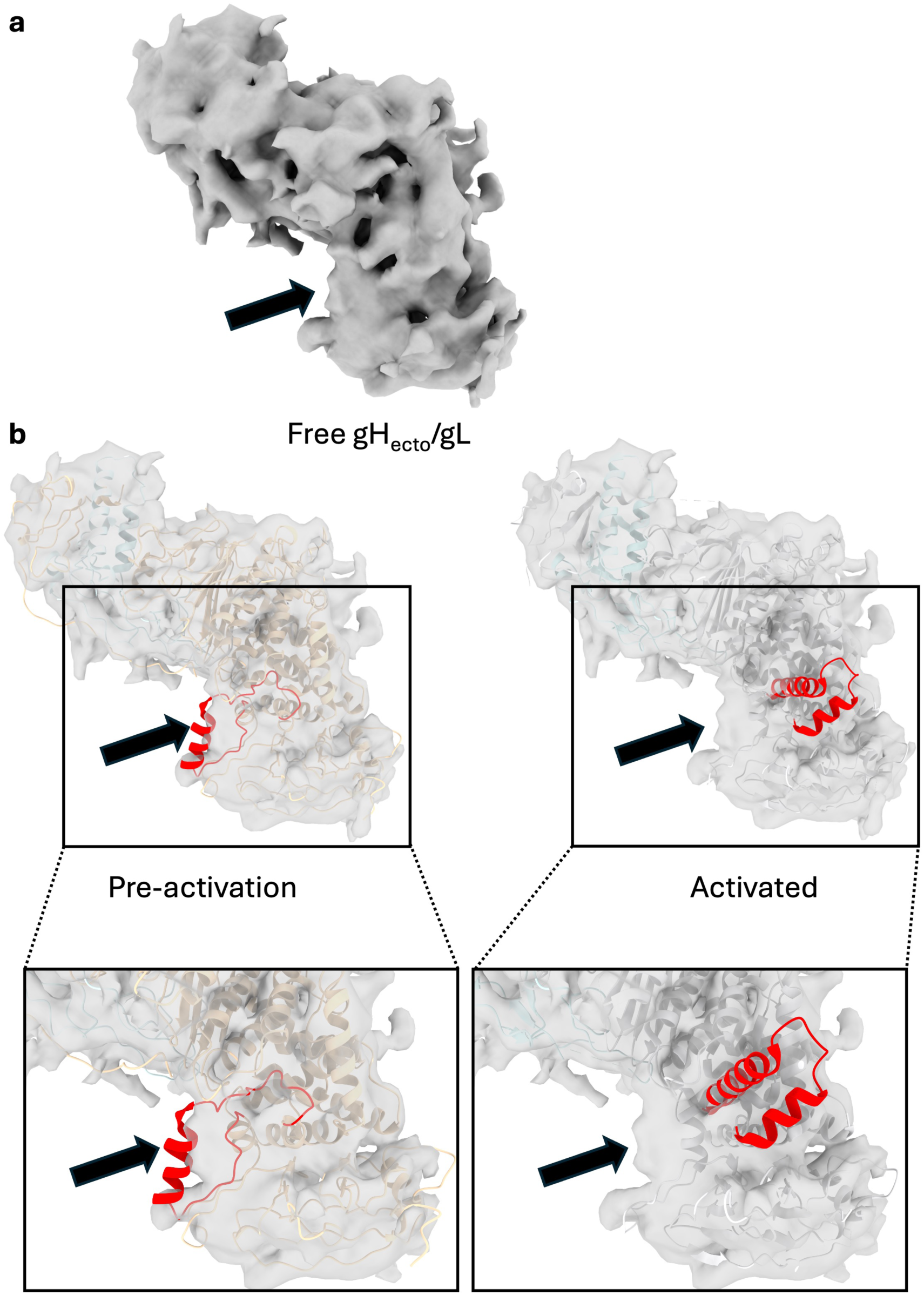
Unbound gH_ecto_/gL more closely resembles CHL27-bound gH_ecto_/gL than gH_ectoΔ48_/gL. **a)** 6.11 angstrom EM reconstruction of unbound gH_ecto_/gL. Arrow points at putative switch density. **b)** ChimeraX fit of pre-activation gH_ecto_/gL (left) and activated gH_ectoΔ48_/gL (right) into free gH_ecto_/gL EM map. Indicated density is empty when gH_ectoΔ48_/gL is fit, whereas switch motif fits reasonably well in the density. Inset zoomed into switch region.

**Extended Data Figure 5.**
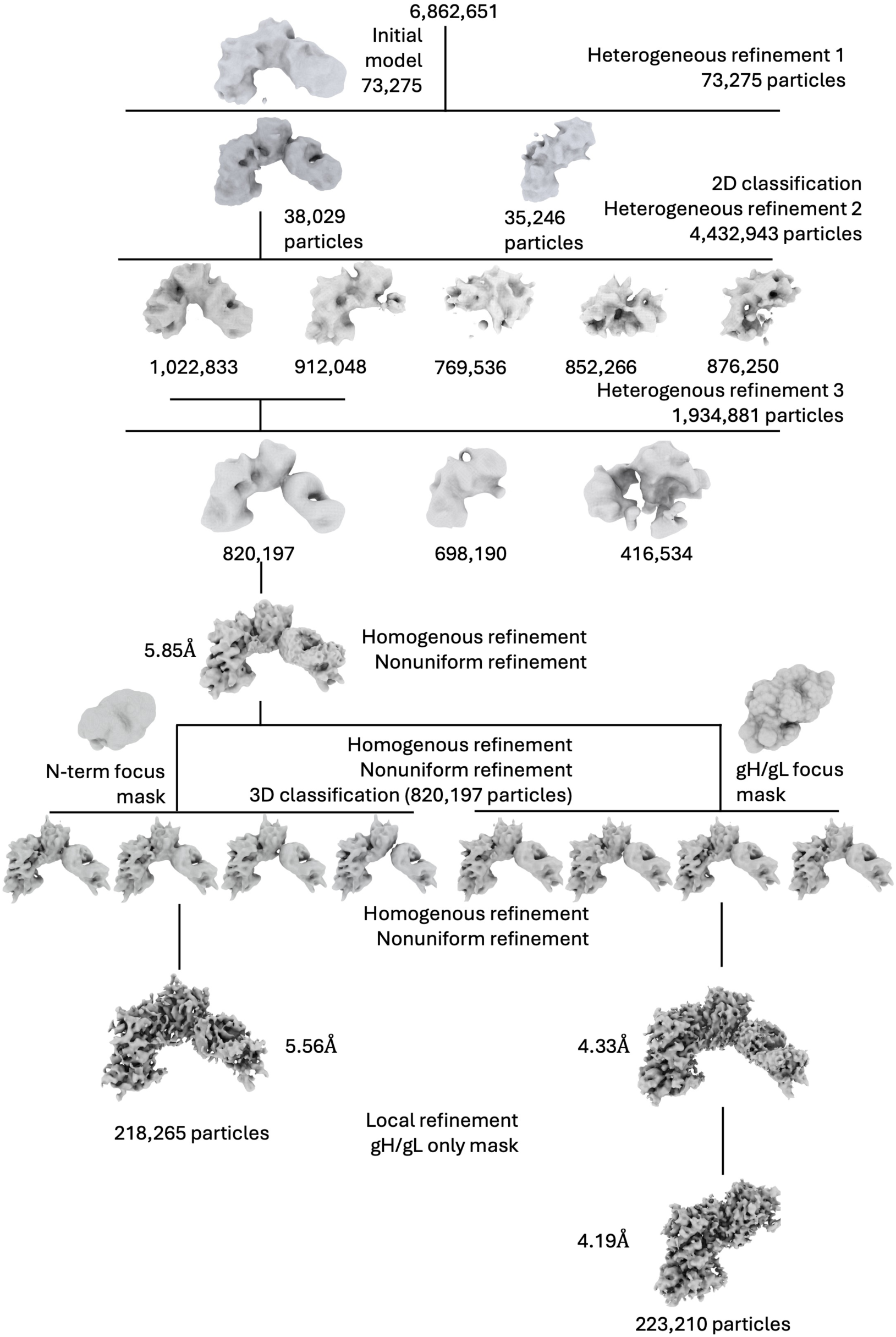
CryoEM analysis workflow.

**Extended Data Figure 6.**
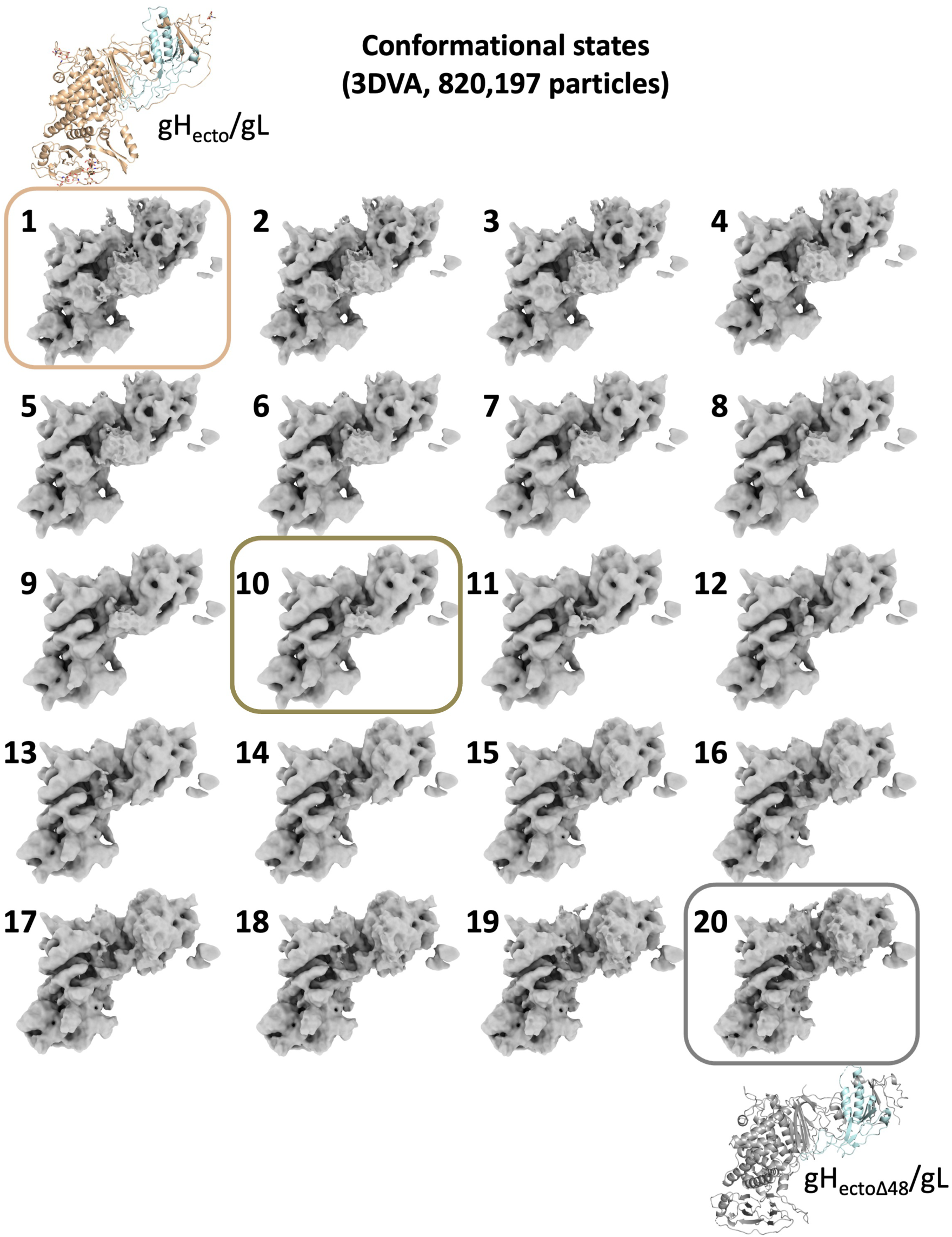
3D Variability analysis-generated maps of gH/gL-focused masking reveal the conformational continuum of gH_ecto_/gL when in complex with CHL27_Fab_. 3DVA resolved continuous heterogeneity across 20 volumes. State 1 (boxed in wheat) refers to the pre-activation conformation, state 10 (boxed in dark brown) refers to an intermediate conformation, and state 20 (boxed in gray) refers to the activated conformation.

**Extended Data Figure 7.**
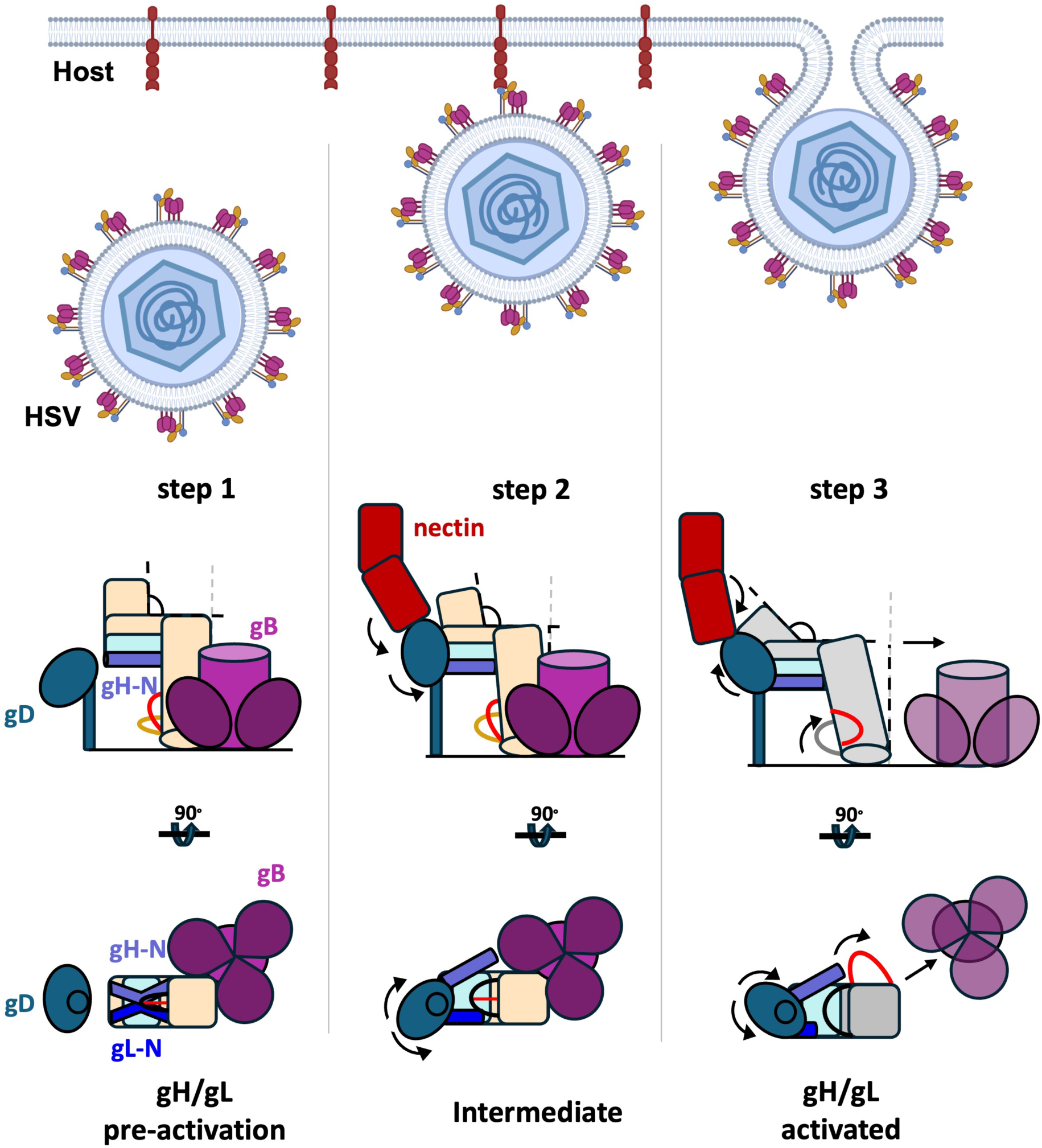
Stepwise “flip-switch” model for allosteric signal transduction through gH/gL. **Step 1:** Prior to host-receptor binding, gH/gL (wheat and pale cyan, respectively) and gB (maroon/dark pink) are locked in a mutually inhibited complex on the virion surface. **Step 2:** host-receptor (nectin, brick) binds gD (sapphire) and leads to gD-gH/gL ectodomain interactions. Movements between nectin, gD, and gH/gL release gL N terminus from its contacts with the gH picket fence. **Step 3:** Release of the N termini leads to a relaxation and tilting of gH/gL that widens the distance between outer β-strands of the C-terminal dock, allowing the switch motif to flip out toward the gB-binding surface of the heterodimer. We posit this destabilizes gH/gL-gB interactions, freeing the labile prefusion gB to transition to its extended intermediate conformation.

**Extended Data Figure 8.**
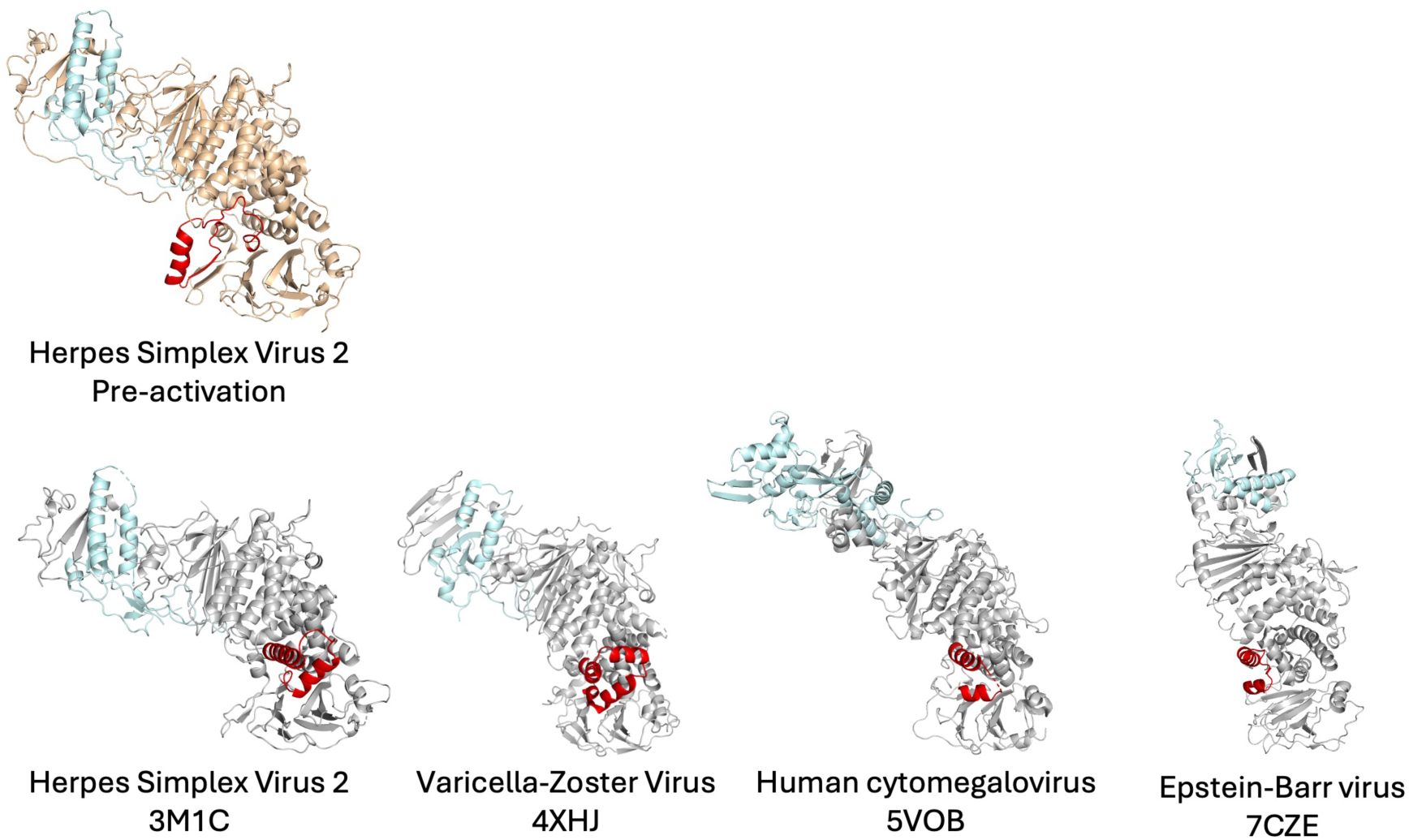
Switch motif is conserved across all known structures of *Herpesviridae* gH/gL homologs. Structures of human-infecting herpesvirus gH/gL ectodomain (PDBIDs: HSV-2 3M1C, VZV 4XHJ, HCMV 5VOB, EBV 7CZE) all show conservation of the switch in its activated, helix-loop-helix configuration. Our structure of HSV-2 gH_ecto_/gL is the only known structure with the switch in its pre-activation conformation. Models were aligned in Pymol by domains H3/D4.

**Extended Data Figure 9.**
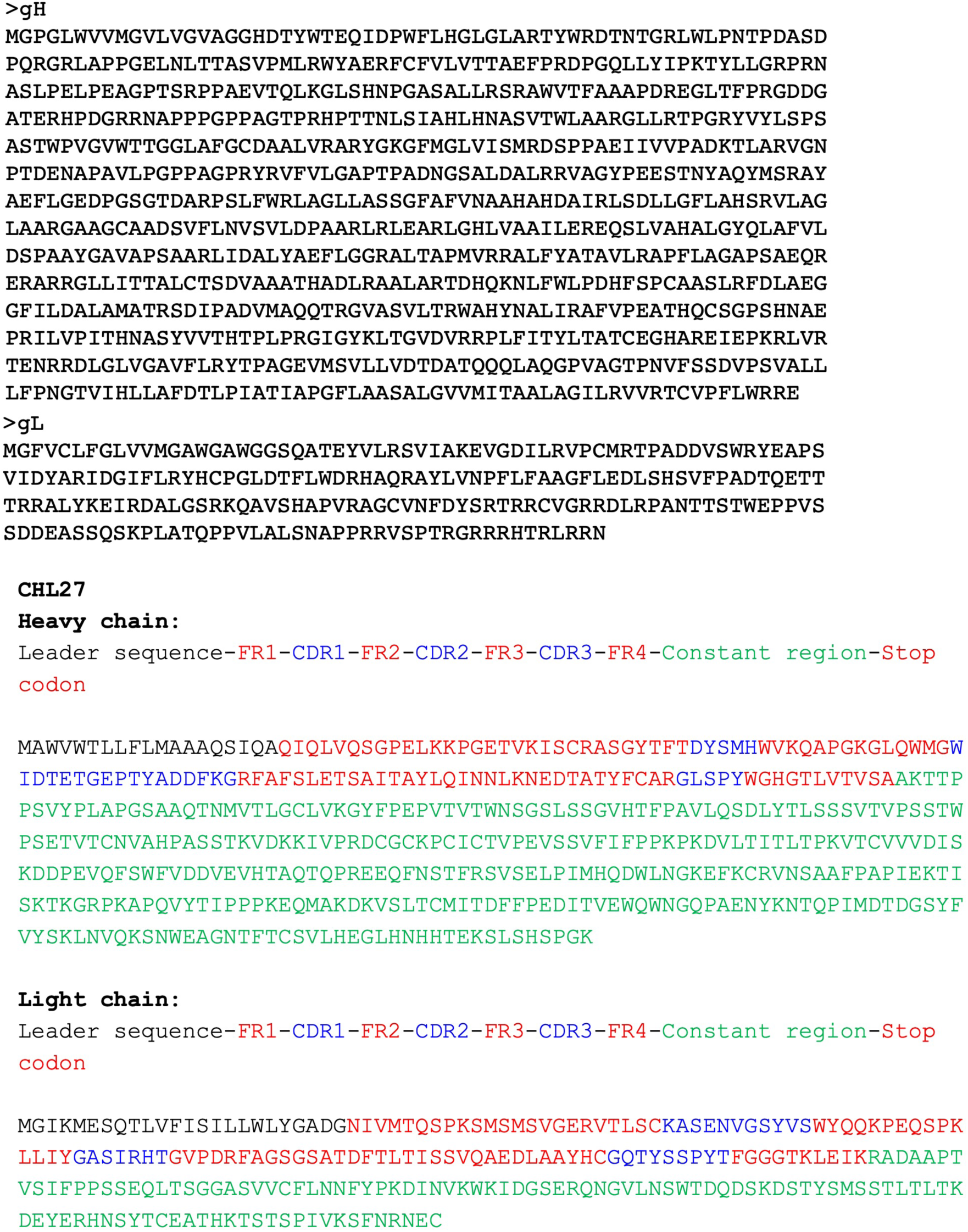
gH/gL and CHL27 sequence and domain maps.

**Extended Data Figure 10.**
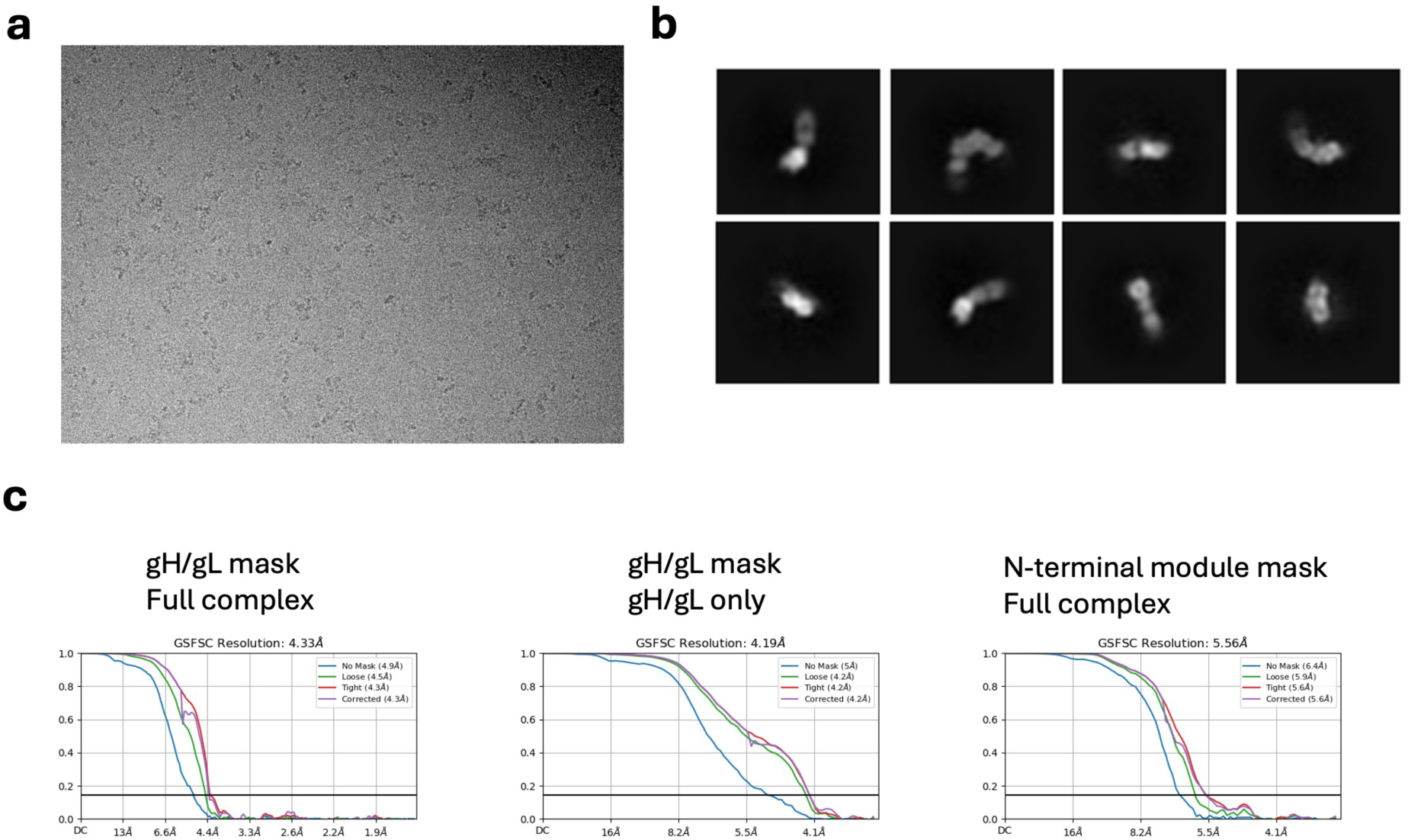
CryoEM data collection and analysis. **a)** Representative cryoelectron micrograph collected on Titan Krios “Caitlin” at the New York Structural Biology Center. **b)** 2D classes for the 820,197 particles before focused 3D classification. **c)** Gold-standard Fourier shell correlation curves for the three reconstructed maps used for model building and structural analysis.

**Extended Data Table 1.** Cryo-EM data collection, refinement, and validation statistics.

| **Data collection and processing** | **gH<sub>ecto</sub>/gL2 only**<br>**PDBID: 9CZG** | **gH<sub>ecto</sub>/gL-CHL27**<br>**PDBID: 9CZR** |
| --- | --- | --- |
| Magnification | 105,000x |  |
| Voltage (kV) | 300 |  |
| Electron exposure (e<sup>-</sup>/ Å<sup>2</sup>) | ~64 |  |
| Defocus range (μm) | -1.2 - -2.2 |  |
| Pixel size (Å) | 0.825 |  |
| Symmetry imposed | C1 |  |
| Initial particle images (no.) | 6,862,651 |  |
| Final particle images (no.) | 223,210 | 218,265 |
| Map resolution (Å) at 0.143 FSC threshold | 4.19 | 5.56 |
| **Refinement** |  |  |
| Initial model used (PDB code) | 3M1C |  |
| Map sharpening *B* factor (Å<sup>2</sup>) |  |  |
| gH/gL local refinement C-term focus | -196 | -256 |
| Composition (no.) |  |  |
| Chains | 2 | 4 |
| Non-hydrogen atoms | 7113 | 8827 |
| Residues | 916 | 1138 |
| Ligands | 7 (BMA:2, NAG: 5) | 7 (BMA:2, NAG: 5) |
| Bonds (RMSD) |  |  |
| Length (Å) (# > 4σ) | 0.008 (14) | 0.103 (71) |
| Angles (°) (# > 4σ) | 1.46 (86) | 2.88 (205) |
| MolProbity score | 2.48 | 2.66 |
| Clash score | 19.5 | 27.4 |
| Ramachandran plot (%) |  |  |
| Outliers | 0.88 | 0.98 |
| Allowed | 3.96 | 4.02 |
| Favored | 95.2 | 95.0 |
| Rotamer outliers (%) | 2.94 | 3.22 |
| Model vs. Data |  |  |
| CC (mask) | 0.77 | 0.58 |
| CC (box) | 0.79 | 0.67 |
| CC (peaks) | 0.68 | 0.49 |
| CC (volume) | 0.77 | 0.59 |
| Mean CC for ligands | 0.72 | 0.70 |

## REFERENCES

1. N., F., Bernard, 1950-, K., David M. (David Mahan), & M., H., Peter. Fields virology. (2013).

2. Atanasiu, D., Saw, W. T., Cohen, G. H. & Eisenberg, R. J. Cascade of Events Governing Cell-Cell Fusion Induced by Herpes Simplex Virus Glycoproteins gD, gH/gL, and gB▿. J Virol 84, 12292–12299 (2010).

3. Liu, L. Fields Virology, 6th Edition. Clin Infect Dis 59, 613–613 (2014).

4. Jeffery-Smith, A. & Riddell, A. Herpesviruses. Medicine 49, 780–784 (2021).

5. Eisenberg, R. J. et al. Herpes Virus Fusion and Entry: A Story with Many Characters. Viruses 4, 800–832 (2012).

6. Poole, C. L. & James, S. H. Antiviral Therapies for Herpesviruses: Current Agents and New Directions. Clin. Ther. 40, 1282–1298 (2018).

7. Heldwein, E. E. & Krummenacher, C. Entry of herpesviruses into mammalian cells. Cell Mol Life Sci 65, 1653–1668 (2008).

8. Pino, G. L. G.-D. & Heldwein, E. E. Well Put Together-A Guide to Accessorizing with the Herpesvirus gH/gL Complexes. Viruses 14, 296 (2022).

9. Connolly, S. A., Jardetzky, T. S. & Longnecker, R. The structural basis of herpesvirus entry. Nat. Rev. Microbiol. 19, 110–121 (2021).

10. Cairns, T. M. & Connolly, S. A. Entry of Alphaherpesviruses. Curr. Issues Mol. Biol. 41, 63–124 (2020).

11. Lu, G. et al. Crystal Structure of Herpes Simplex Virus 2 gD Bound to Nectin-1 Reveals a Conserved Mode of Receptor Recognition. J Virol 88, 13678–13688 (2014).

12. Giovine, P. D. et al. Structure of Herpes Simplex Virus Glycoprotein D Bound to the Human Receptor Nectin-1. Plos Pathog 7, e1002277 (2011).

13. Connolly, S. A. et al. Structure-Based Analysis of the Herpes Simplex Virus Glycoprotein D Binding Site Present on Herpesvirus Entry Mediator HveA (HVEM). J Virol 76, 10894–10904 (2002).

14. Stiles, K. M. et al. Herpes Simplex Virus Glycoprotein D Interferes with Binding of Herpesvirus Entry Mediator to Its Ligands through Downregulation and Direct Competition▿. J Virol 84, 11646–11660 (2010).

15. Shukla, D. et al. A Novel Role for 3-O-Sulfated Heparan Sulfate in Herpes Simplex Virus 1 Entry. Cell 99, 13–22 (1999).

16. Tiwari, V. et al. Soluble 3-O-sulfated heparan sulfate can trigger herpes simplex virus type 1 entry into resistant Chinese hamster ovary (CHO-K1) cells. J Gen Virol 88, 1075–1079 (2007).

17. White, J. M., Ward, A. E., Odongo, L. & Tamm, L. K. Viral Membrane Fusion: A Dance Between Proteins and Lipids. Annu. Rev. Virol. 10, 139–161 (2023).

18. Harrison, S. C. Viral membrane fusion. Virology 479-480C, 498–507 (2015).

19. Chowdary, T. K. et al. Crystal structure of the conserved herpesvirus fusion regulator complex gH-gL. Nat. Struct. Mol. Biol. 17, 882–8 (2010).

20. Cairns, T. M. et al. Capturing the Herpes Simplex Virus Core Fusion Complex (gB-gH/gL) in an Acidic Environment. J Virol 85, 6175–6184 (2011).

21. Cairns, T. M. et al. Localization of the Interaction Site of Herpes Simplex Virus Glycoprotein D (gD) on the Membrane Fusion Regulator, gH/gL. J Virol 94, (2020).

22. Pataki, Z., Viveros, A. R. & Heldwein, E. E. Herpes Simplex Virus 1 Entry Glycoproteins Form Complexes before and during Membrane Fusion. MBio 13, e0203922 (2022).

23. Atanasiu, D. et al. Regulation of herpes simplex virus gB-induced cell-cell fusion by mutant forms of gH/gL in the absence of gD and cellular receptors. MBio 4, e00046–13 (2013).

24. Atanasiu, D., Saw, W. T., Cairns, T. M., Eisenberg, R. J. & Cohen, G. H. Using Split Luciferase Assay and Anti-Herpes Simplex Virus Glycoprotein Monoclonal Antibodies To Predict a Functional Binding Site between gD and gH/gL. J Virol 95, (2021).

25. Cairns, T. M., Landsburg, D. J., Whitbeck, J. C., Eisenberg, R. J. & Cohen, G. H. Contribution of cysteine residues to the structure and function of herpes simplex virus gH/gL. Virology 332, 550–562 (2005).

26. Punjani, A. & Fleet, D. J. 3DFlex: determining structure and motion of flexible proteins from cryo-EM. Nat. Methods 20, 860–870 (2023).

27. Punjani, A. & Fleet, D. J. 3D variability analysis: Resolving continuous flexibility and discrete heterogeneity from single particle cryo-EM. J. Struct. Biol. 213, 107702 (2021).

28. Zhong, E. D., Bepler, T., Berger, B. & Davis, J. H. CryoDRGN: reconstruction of heterogeneous cryo-EM structures using neural networks. Nat. Methods 18, 176–185 (2021).

29. Walsh, R. M. et al. Structural principles of distinct assemblies of the human α4β2 nicotinic receptor. Nature 557, 261–265 (2018).

30. Si, Z. et al. Different functional states of fusion protein gB revealed on human cytomegalovirus by cryo electron tomography with Volta phase plate. Plos Pathog 14, e1007452 (2018).

31. Vanarsdall, A. L., Howard, P. W., Wisner, T. W. & Johnson, D. C. Human Cytomegalovirus gH/gL Forms a Stable Complex with the Fusion Protein gB in Virions. Plos Pathog 12, e1005564 (2016).

32. Cooper, R. S., Georgieva, E. R., Borbat, P. P., Freed, J. H. & Heldwein, E. E. Structural basis for membrane anchoring and fusion regulation of the Herpes Simplex Virus fusogen gB. Nat. Struct. Mol. Biol. 25, 416–424 (2018).

33. Pataki, Z., Sanders, E. K. & Heldwein, E. E. A surface pocket in the cytoplasmic domain of the herpes simplex virus fusogen gB controls membrane fusion. PLOS Pathog. 18, e1010435 (2022).

34. Hannah, B. P. et al. Herpes Simplex Virus Glycoprotein B Associates with Target Membranes via Its Fusion Loops. J. Virol. 83, 6825–6836 (2009).

35. White, E. M., Stampfer, S. D. & Heldwein, E. E. Herpes Simplex Virus, Methods and Protocols. Methods Mol Biology 2060, 377–393 (2019).

36. Zheng, S. Q. et al. MotionCor2: anisotropic correction of beam-induced motion for improved cryo-electron microscopy. Nat. Methods 14, 331–332 (2017).

37. Rohou, A. & Grigorieff, N. CTFFIND4: Fast and accurate defocus estimation from electron micrographs. J. Struct. Biol. 192, 216–221 (2015).

38. Punjani, A., Rubinstein, J. L., Fleet, D. J. & Brubaker, M. A. cryoSPARC: algorithms for rapid unsupervised cryo-EM structure determination. Nat. Methods 14, 290–296 (2017).

39. Fernandez-Leiro, R. & Scheres, S. H. W. A pipeline approach to single-particle processing in RELION. Acta Crystallogr. Sect. D 73, 496–502 (2017).

40. Goddard, T. D. et al. UCSF ChimeraX: Meeting modern challenges in visualization and analysis. Protein Sci. 27, 14–25 (2018).

41. Emsley, P., Lohkamp, B., Scott, W. G. & Cowtan, K. Features and development of Coot. Acta Crystallogr Sect D Biological Crystallogr 66, 486–501 (2010).

42. Croll, T. I. ISOLDE: a physically realistic environment for model building into low-resolution electron-density maps. Acta Crystallogr. Sect. D: Struct. Biol. 74, 519–530 (2018).

43. Adams, P. D. et al. PHENIX: a comprehensive Python-based system for macromolecular structure solution. Acta Crystallogr Sect D Biological Crystallogr 66, 213–221 (2010).

44. Lyskov, S. et al. Serverification of Molecular Modeling Applications: The Rosetta Online Server That Includes Everyone (ROSIE). PLoS ONE 8, e63906 (2013).

