## Supplementary material for "Allosteric mechanism of membrane fusion activation in a herpesvirus": Legends for Supplemental Movies

**The PDF file includes:**

Legends for Supplemental Movies S1 to S6

**Supplemental Movie S1. Conformational rearrangements between pre-activation and activated gH/gL states.** Morph generated in Pymol between the $gH_{ecto}$/gL structure and the $gH_{ecto\Delta48}$/gL structure (PDBID: 3M1C) showing the rearrangements in the model of the gH/gL ectodomain as it transitions from the pre-activation to activated state.

**Supplemental Movie S2. Face-on view of volume series generated by cryoSPARC's 3D Flexibility analysis showing large-scale movements of $gH_{ecto}$/gL-CHL27$_{Fab}$ complex.**

**Supplemental Movie S3. Top-down view of volume series generated by cryoSPARC's 3D Flexibility analysis showing large-scale movements of $gH_{ecto}$/gL-CHL27$_{Fab}$ complex.**

**Supplemental Movie S4. Volume series generated by cryoSPARC's 3D Variability analysis showing local conformational rearrangements of the gL N terminus and the C-terminal switch motif in $gH_{ecto}$/gL.** Movie view is facing the gL N terminus.

**Supplemental Movie S5. Volume series generated by cryoSPARC's 3D Variability analysis showing local conformational rearrangements of the gL N terminus and the C-terminal switch motif in $gH_{ecto}$/gL.** Movie S5 view is rotated 180 degrees from movie S4 to face the gH N terminus and switch region.

**Supplemental Movie S6. Volume series generated by cryoDRGN on the same particle set validates cryoSPARC's 3DFlex and 3DVariability analyses.** Translation of the gL N terminus across the N-terminal module and concurrent disappearance of the helical switch density recapitulates CryoSPARC's 3DVA-generated volume series.
